# Efficient Enumeration and Visualization of Helix-Coil Ensembles

**DOI:** 10.1101/2023.09.16.558052

**Authors:** Roy Hughes, Shiwen Zhao, Terrence G. Oas, Scott C. Schmidler

**Affiliations:** Department of Biochemistry, Duke University, Durham, North Carolina; Program in Computational Biology and Bioinformatics, Duke University, Durham, North Carolina; Department of Statistical Science, Duke University, Durham, North Carolina

## Abstract

Helix-coil models are routinely used to interpret CD data of helical peptides or predict the helicity of naturally-occurring and designed polypeptides. However, a helix-coil model contains significantly more information than mean helicity alone, as it defines the entire ensemble - the equilibrium population of every possible helix-coil configuration - for a given sequence. Many desirable quantities of this ensemble are either not obtained as ensemble averages, or are not available using standard helicity-averaging calculations. Enumeration of the entire ensemble can allow calculation of a wider set of ensemble properties, but the exponential size of the configuration space typically renders this intractable. We present an algorithm that efficiently approximates the helix-coil ensemble to arbitrary accuracy, by sequentially generating a list of the *M* highest populated configurations in descending order of population. Truncating this list of (configuration, population) pairs at a desired accuracy provides an approximating sub-ensemble. We demonstrate several uses of this approach for providing insight into helix-coil ensembles and folding mechanisms, including landscape visualization.

**SIGNIFICANCE**
Helix-coil models define the probability distribution of helix-coil configurations for a polypeptide (a helix-coil ensemble). Each configuration specifies which residues are α-helical and which are not. We used an accurate helix-coil model, paired with concepts from the field of probabilistic graphical modeling, to devise an algorithm capable of enumerating helix-coil configurations in order of decreasing probability. By enumerating ensembles for a representative set of peptides we find that helix-coil ensembles tend to be highly concentrated, with the vast majority of probability mass assigned to a relatively small set of configurations from a configuration space that is often astronomical in size. This result facilitates the development of new and accurate methods for analyzing and predicting the behavior of helical polypeptides.

## INTRODUCTION

Helix-coil theory (1) is a framework developed in the 1950’s to describe the transition of biopolymers between the conformational state consistent with the α-helix of Linus Pauling (2) and the disordered random coil state described in traditional polymer theory (3). Important statistical mechanical models developed from helix-coil theory in the 1950’s and 1960’s, called *helix-coil models*, include the Zimm-Bragg (4) and Lifson-Roig (5) models. Advances in polypeptide synthesis technology (6) in the 1980’s allowed researchers to easily construct experimental model systems for studying the helix-coil transition within short peptides. Experiments on these peptide model systems greatly expanded the set of helicity observations recorded in the literature. In many cases helix-coil models were used to provide a biophysical interpretation for the observations obtained. These studies in turn led to numerous refinements and extensions to the Lifson-Roig and Zimm-Bragg models to account for unexplained variances present in the new data sets (7–15). A substantive result of these lines of research effort has been the development of accurate, predictive helix-coil models (10, 16–18) that have set the standard by which other protein structure prediction algorithms are often judged (19–21). At present, helix-coil models are arguably the most accurate predictive models of polypeptide structure available when judged by their consistency with experimental measurements.

A helix-coil model defines a *helix-coil ensemble* associated with a specified polypeptide sequence and its dependence on temperature, pH, and other solvent conditions. A helix-coil ensemble is a probability distribution defined over a space of helix-coil configurations (conformations) for a given polypeptide. Each configuration specifies whether a given residue adopts backbone (*ϕ, ψ*) angles consistent with those found within an *α*-helix (helical residue) or not (coil residue). For a polypeptide of *N* residues there are 2^*N*^ possible helix-coil configurations, usually represented as a string of h and c characters, which respectively indicate helical or coil residues (e.g. ccchhcch). While a helix-coil ensemble assigns a probability to each configuration in this space, nearly all experimental studies of helical peptides by necessity measure only ensemble average properties. The most important of these observables is the mean chain helicity, which is easily measured using circular dichroism (22). This observable is the primary means by which the parameters of helix-coil models have been estimated and their predictive accuracy evaluated(16, 18). Fortunately efficient algorithms exist for calculating many ensemble-averaged observable properties(18) such as mean helicity. Other commonly computed ensemble average observables include the probability that a specific residue is helical (residue-specific helicity) and the probability that an amide proton of residue *k* participates in a hydrogen bond with the carbonyl oxygen of residue *k*−4 (the fractional amide protection).

However, averaging can hide important features of an ensemble and fail to provide a complete characterization of the system. For example, if the ensemble is multimodal (see e.g Figure 4), then the mean is a poor summary. Yet examination of the entire ensemble is inhibited by the exponentially large size of the configuration space: for a 27 residue polypeptide the ensemble comprises over 134 million distinct configurations. This calculation, however, ignores the fact that different configurations have widely varying probabilities of occurrence. In fact, a (relatively) small number of configurations may constitute the bulk of the populated space in a given ensemble and dominate contributions to the partition function. This suggests the possibility of accurately approximating an ensemble with a much smaller set of configurations. The problem considered in this paper is to enumerate the configuration space *in order of descending probability of occurrence*, allowing the construction of highest probability configuration sets. These sets constitute (1 − *ϵ*) -approximate ensembles, providing accuracy up to a user-defined tolerance E. (Alternatively, our approach can be used to find the approximate ensemble of pre-specified size *M* which achieves the minimal approximation error *ϵ*^*^, along with determining the corresponding value of ϵ^*^.) This enables, among other things, the calculation of a (1−*ϵ*) -approximation for the partition function, a problem that has been studied in other contexts (23, 24).

In what follows we provide an efficient algorithm for solving this ensemble approximation problem, which leverages the connection between helix-coil models and probablistic graphical models noted previously(18). With this in hand, we then explore some ways in which approximate ensemble enumeration can be used to gain new insights about helix-coil ensembles. Analyzing a large database of synthetic and naturally-occurring helical peptides, we find that most helix-coil ensembles are highly concentrated, populating only a small fraction of the allowed configuration space. We also use ensemble enumeration to understand the errors arising from the “single-sequence approximation” used previously by some authors. Finally, we demonstrate the use of ensemble approximation to aid in visualization of entire helix-coil ensembles, and provide examples of how this may be a powerful tool for researchers to improve understanding of helix-coil systems.

## METHODS

### The Helix Coil Model

The work in this study was performed using a helix-coil model developed by Schmidler et al. (18), which was fit to the extensive data reported by Baldwin, Kallenbach and others(7–9, 25–28). To a large extent the form of the energy function of the model is adapted from previous work by Serrano and co-workers (10, 15, 16), which was in turn inspired by the helix-coil models used to interpret the experimental data (7–9, 28). The parameter values used here were determined by statistical fitting to a large (1397 data points) peptide helicity database using Bayesian inference. The resulting model has a high predictive accuracy(18). Here we briefly introduce some notation and summarize the model.

Let Ω = (*ω*_1_, …, *ω*_*l*_) denote the sequence of a peptide of length *l*, with each ω_*i*_ taking values in the set of twenty amino acids. Let *X* = (*x*_1_,, *x*_*l*_)be a binary vector representing a corresponding helix-coil configuration, with *x*_*i*_ = 1 if residue *i* is helical and *x*_*i*_ = 0 indicating coil. A given peptide may or may not have terminal blocking groups attached; if there is no N-terminal blocking group (typically acetyl) then we have the constraint *x*_1_ = 0 and if there is no C-terminal blocking group (typically amide) then *x*_*l*_ = 0. Let *b*_*N*_ = 1 if the N-terminus of the peptide is blocked and *b*_*N*_ = 0 if it is not blocked; likewise, *b*_*C*_ = 1 if the C-terminus of the peptide is blocked and *b*_*C*_ = 0 if unblocked. Denote by *χ* the set of possible configurations; then the number of allowed configurations |*χ*| is 2^*l*−2^, 2^*l*−1^, or 2^*l*^ when there are zero, one, or two blocked termini, respectively. The energy of a configuration *X*, relative to the reference (all-coil) configuration, is given by

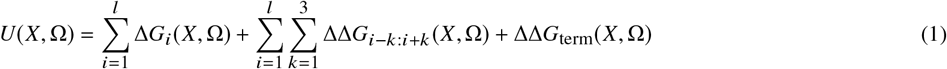

where Δ*G*_*i*_ is the contribution of residue *i* to the free energy relative to the reference state, and ΔΔ*G*_*i*−*k,i*+*k*_ contains additional energetic contributions from sidechain-sidechain and sidechain-backbone interactions spanning the corresponding window of residues (of length 2*k* + 1) centered at sequence position *i*. Details of the energetic model are given in the supplementary material. The probability (or population) of a given configuration *X* is given by

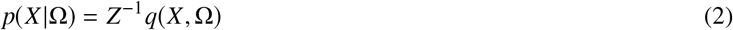

where *q*(*X*, Ω) = *e*^−*βU*(*X*,Ω)^ is the “statistical weight” of configuration *X, β* = (*RT*)^−1^ for solvent temperature *T* and ideal gas constant *R*, and *Z* = Σ_*X* ∈ *χ*_ *q*(*X*, Ω) is the partition function. Schmidler et. al (18) provided an efficient algorithm for exact calculation of *Z*, as well as of the ensemble averaged chain helicity

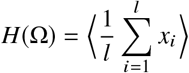

Here ⟨ ·⟩denotes expectation with respect to *p* (*X*| Ω). The parameter values used in this paper have been estimated using Bayesian inference from a large database of peptide helicity measurements as described previously(18).

In the historical development of helix-coil theory, the statistical weight is often ascribed to residue-specific contributions 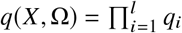, with *q*_*i*_ being the statistical weight of the *i*^th^ residue. We can write our model in this form by defining:

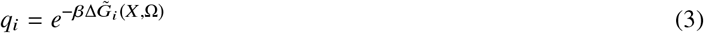

with the Δ*G* terms redefined to incorporate the ΔΔ*G* interactions as well:

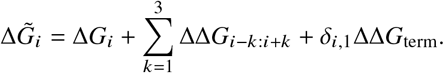

We will find it convenient to adopt this parameterization of the model in the next section.

### Efficient Ensemble Enumeration

In this section we provide an algorithm for enumerating the *M* highest probability configurations in a helix-coil ensemble in order of descending probability. We call such a set of *M* configurations a *sub-ensemble*. When *M* is chosen as 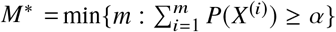, we call the sub-ensemble a (1 − α)*-approximate ensemble*. We show that the helix-coil model can be represented as a probabilistic graphical model, and exploit this representation to provide an algorithm for enumerating sub-ensembles efficiently. In particular, the algorithm provided is *O* (*l*^2^*M*\*), so if *M*^*^ grows at most polynomially in *l*, then the approximation algorithm is fully polynomial in *l*.

#### Ensemble enumeration

We define a helix-coil *ensemble* Ψ of a peptide as a sequence of *N* ordered pairs

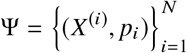

Here *X*^(*i*)^ is the *i*th most probable helix-coil configuration, *p*_*i*_ is its probability (so *i* is the *rank* of *X*^(*i*)^), and *N* = |Ψ| is the size of the helix-coil configuration space, given by

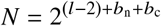

where *b*_n_ (resp. *b*_c_) is a binary indicator taking value 1 if an N-terminal (resp. C-terminal) blocking group is present, and 0 otherwise. For a given ensemble Ψ we define

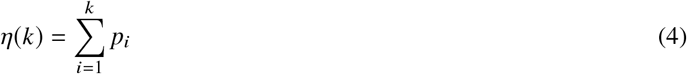

to be the cumulative distribution function (cdf) of *p*_*i*_. Then η(*k*) gives the proportion of the ensemble Ψ that is accounted for by the *k* most probable configurations. We define the (generalized) inverse-cdf η^−1^(α)

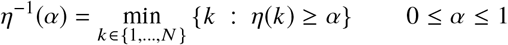

which gives the *sub-ensemble size corresponding to proportion* α *of* Ψ. That is, η^−1^ α is the size of the minimal set of configurations within Ψ that have a total population greater than or equal to α. We define Ψ_*α*_ as the *minimal sub-ensemble of* Ψ *corresponding to population* α:

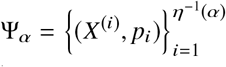

Thus Ψ_*α*_ is the sub-ensemble consisting of the *η*^−1^(*α*) most probable configurations in Ψ, with Ψ_1_ = Ψ. We will define an enumeration algorithm that takes as input some *α* ∈ (0, 1), and outputs Ψ_*α*_ as a (1 − α)-approximation of the full ensemble Ψ.

### The Helix-coil Model as a Probabilistic Graphical Model

A probabilistic graphical model (PGM) is a representation of a stochastic model using a marriage of probability theory and graph theory. PGMs provide a compact representation of models, along with efficient computational algorithms for performing probabilistic inference when the model satisfies certain conditions on its conditional independence structure.(29, 30). PGMs are used extensively in a variety of fields including machine learning(31) and biology(32).

We will use a particular form of PGM known as an *undirected graphical model* (UGM) or *Markov random field*. A UGM consists of a *graph G* = (*V, E*) made up of vertices *V* and edges *E* ⊂ *V* × *V*, along with a set of non-negative potential functions 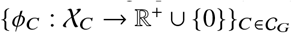 defined on the cliques of *G*. (A *clique C* ∈ C_*G*_ is a set of vertices in a graph *G* whose induced subgraph is *complete*, or fully connected; χ_*C*_ denotes the set of instantiations of the variables in *C*.) Vertices in the graph represent random variables, and edges connecting the vertices represent direct dependency relationships (energetic interactions). The PGM defines a joint distribution over all variables in the graph which is proportional to the product of the potential functions:

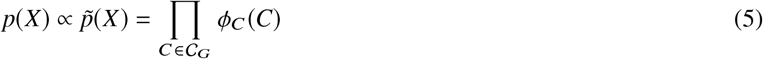

where C_*G*_ is the set of cliques in *G*.

In a helix-coil model, the random variables are the residues in the polypeptide sequence Ω =(*ω*_1_, …, *ω*_*l*_)and the unobserved configuration indicators *X* =(*x*_1_, …, *x*_*l*_). As described previously, each residue *ω*_*i*_ has a relative free energy that depends on its configuration *x*_*i*_ and the identities and configurations of other residues that surround it. The vertex associated with a random variable (*x*_*i*_ or *ω*_*i*_) is connected by an edge to the vertices of every other residue or configuration indicator on which the relative free energy of residue *i* depends. A visual depiction of the graph for an eight residue peptide is given in Figure 1.

**Figure 1.**
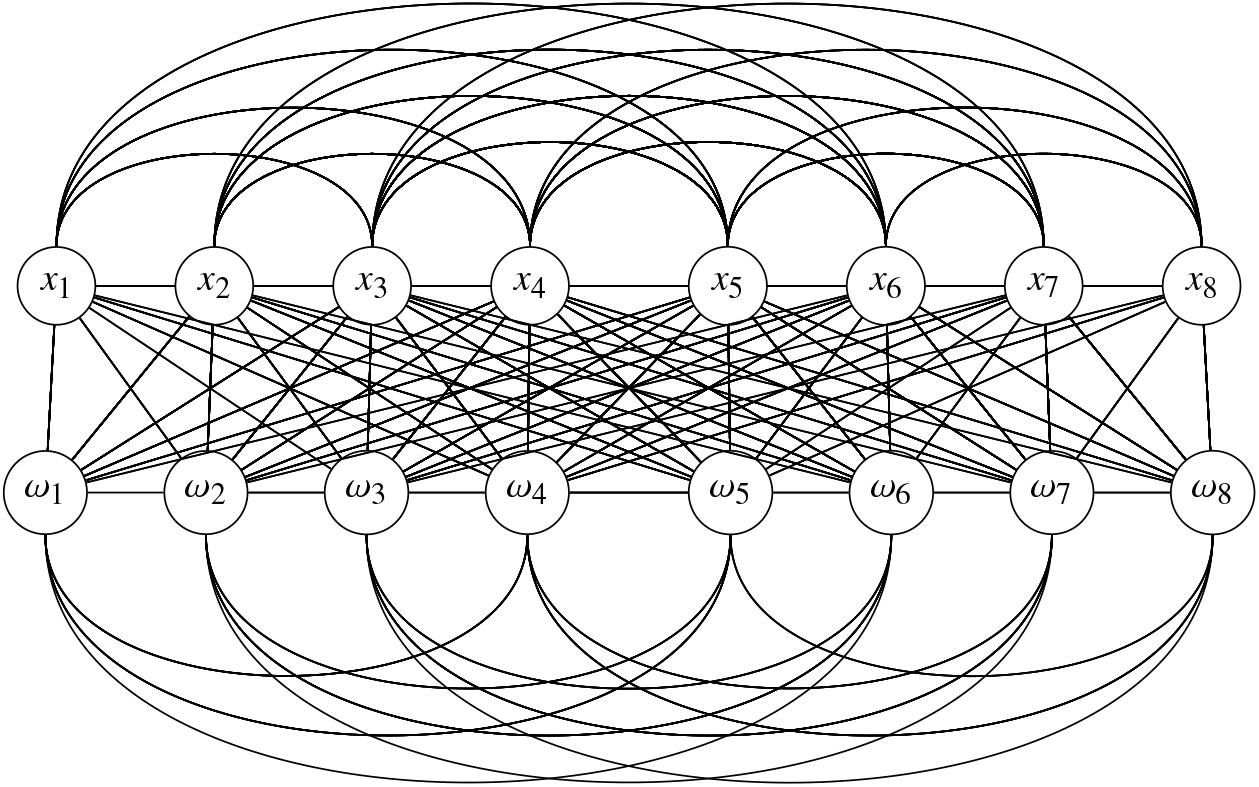
Undirected graphical model representation of the helix-coil model for an 8-residue peptide. Two vertices are connected if and only if they appear together in a product within a term of *U*(*X*, Ω) (1).

The graph in Figure 1 allows direct visual inspection of the energetic interactions in the model, but is somewhat complex. We can simplify the graph by treating the sequence Ω as known (observed). The PGM for the corresponding conditional distribution *p(X* | Ω) from (2) is given in Figure 2, where the observed *ω*_*i*_ values are absorbed into the potentials. In what follows we suppress the explicit conditioning on Ω in (5) to simplify notation.

**Figure 2.**
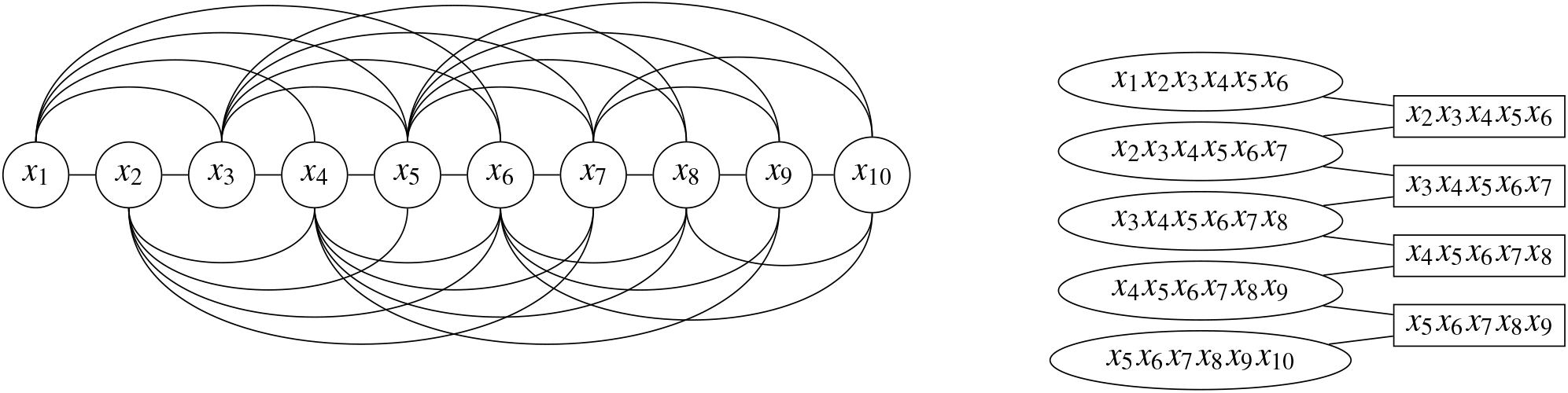
(a) Undirected graphical model representation of the helix coil model for a ten residue peptide when Ω is assumed known. Vertices represent random variables describing the helix/coil conformation of individual sequence positions. Edges denote probabilistic dependencies arising from interaction energies in the model. (b) The clique tree corresponding to the graphical model in (a).

### The Message Passing Algorithm

An important tool for defining the enumeration algorithm is the *message passing algorithm*, a procedure defined on probabilistic graphical models which enables calculation of the partition function, finding the maximum statistical weight configuration, and calculation of marginal probabilities (30).

### Clique tree representation

The message-passing algorithm for ensemble enumeration operates on a transformed graph known as a *clique tree*. Algorithms for generating clique trees from general PGMs are available(30), but the helix coil model gives rise to a particularly simple *linear chain* graph, shown in Figure 2. In a clique tree, each vertex represents a subset of variables in the original graph, and each edge represents the overlapping scope of its two neighboring vertices, known as a *sepset*. In the clique tree for the helix-coil model (Figure 2), every set of six consecutive residues is grouped to form a vertex (clique). For example, the first clique *C*_1_ = {*x*_1_, …, *x*_6_} comprises residues 1 through 6. The model can then be factorized by assigning a potential function to each clique and each sepset, with the unnormalized joint distribution 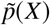 expressed as

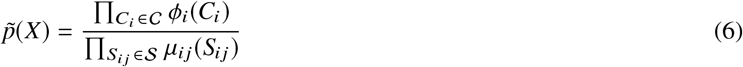

where 𝒮 is the set of sepsets in the clique tree. For the linear graph of the helix-coil tree in Figure 2 this can be written in the slightly simpler form:

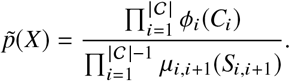

This representation of the unnormalized joint distribution can be achieved by assigning each potential term from (5) to any single clique in the clique tree which contains all *x*_*i*_’s on which the term depends, and setting μ_*i,i*+1_≡1 for *i* = 1, …, *l*−1. A complete set of corresponding clique potentials can be written simply in terms of the *q*_*i*_’s (3):

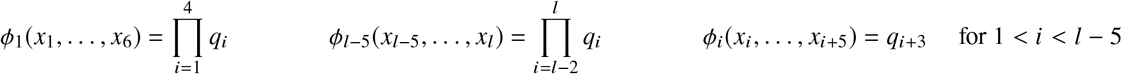

It is straightforward to check that (5) is then equal to *q* (*X*, Ω) in (2). Initializing all sepset potentials to one, (μ_*i,i*+1_≡1) ensures that the property (6) holds.

A key property of the message passing algorithm discussed below, which will modify the ϕ_*i*_’s and μ_*i,j*_’s, is that this property (6) is maintained, i.e. it is *invariant* under message passing operations.

### Message passing

Consider the clique tree in Figure 2. (Although the “tree” here is linear, we adopt the common terminology and designate the first clique *C*_1_ = {*x*_1_, *x*_2_, *x*_3_, *x*_4_, *x*_5_, *x*_6_} as a *leaf* clique, and the last clique *C*_*l*−5_ = {*x*_5_, *x*_6_, *x*_7_, *x*_8_, *x*_9_, *x*_10_} as the *root* clique. The algorithm starts from the clique *C*_1_ and associated potential ϕ_1_(*x*_1_, *x*_2_, *x*_3_, *x*_4_, *x*_5_, *x*_6_), and marginalizes out (sums over) *x*_1_ to generate a new potential δ_1,2_(*x*_2_, *x*_3_, *x*_4_, *x*_5_, *x*_6_) =Σ_*x*1_ ϕ_1_(*x*_1_, *x*_2_, *x*_3_, *x*_4_, *x*_5_, *x*_6_) with scope {*x*_2_, *x*_3_, *x*_4_, *x*_5_, *x*_6_} known as a *sum-flow message*. (If marginalization is replaced with maximization, it is called a *max-flow* message.) The generated message is passed to the second clique where it is multiplied by the second clique’s potential ϕ_2_(*x*_2_, *x*_3_, *x*_4_, *x*_5_, *x*_6_, *x*_7_) to generate an updated potential for the second clique 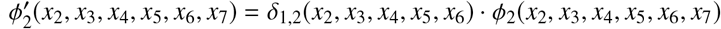. This procedure is repeated along the clique tree until the last (root) clique receives the message from its neighbor. After the root receives the messages and updates its potential, it passes a new message back to its neighbors. The message passing continues until the leaf clique has received a message from it’s neighbor. After these two (left and right, or forward and backward) waves of message passing, the clique tree has reached equilibrium and is referred to as a *calibrated* clique tree. If max-flow messages are used in the message passing, we refer to it as a *max-calibrated* tree.

Denote the new potential associated with each clique following calibration by γ_*i*_. Each sepset is associated with two messages, one for each direction. Denote these two messages as δ_*i,i* +1_ and δ_*i* +1,*i*_, where *i* and *i* + 1 are the two cliques connected by this sepset, and define new potential μ_*i,i* +1_ = δ_*i,i* +1_ · δ_*i* +1,*i*_ on the sepset as the product of these messages. These potentials {γ_*i*_} and {μ_*i,i* +1_} on the calibrated clique tree then exhibit the following properties: (30)

- For a max-calibrated clique tree, 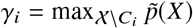 and 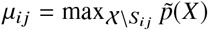, where *C*_*i*_ is the scope of clique *i, S*_*ij*_ is the scope of the sepset connecting cliques *i* and *j*, and 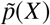 is the unnormalized joint distribution (5). For a sum-calibrated clique tree, these statements hold when substituting max with Σ (i.e. γ_*i*_ gives the correct marginal probability for *C*_*i*_.)
- *Consistency*: 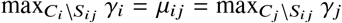 for a max-calibrated tree. For a sum calibrated tree, replace max by Σ.
- The unnormalized joint distribution 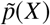 can be expressed as

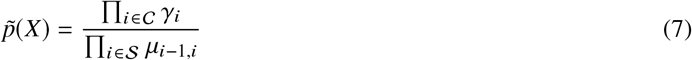

i.e. (6) is left invariant.

### Fast Enumeration of Conformers in Statistical Weight Order

We have seen that after the clique tree is max calibrated, the potential associated with a clique (or sepset) is the marginally maximum function of *p*∼ *X* with respect to the scope of the factor (or sepset). To find the most probable configuration on the calibrated tree we can use the max consistency property defined above. Starting from a leaf clique, find the maximum configuration on that clique. Its neighbor clique must then agree with this maximum configuration over the variables on their shared sepset. Fixing these variables at their maximum configuration values, we can then find the maximum configuration for the remaining variables of the neighboring clique. This procedure is repeated along the clique tree until the entire maximum configuration is determined. In this way, given the max-calibrated clique tree of the probabilistic graphical representation of the helix-coil model, the most populated configuration can be obtained.

However, this provides only the single most probable configuration. Nilsson (33) provided an algorithm to efficiently enumerate the *M* most probable configurations of a probabilistic graphical model, which uses message passing. Let {*X*^(1)^, …, *X*^(*M*)^} be the *M* most probable configurations. An outline of Nilsson’s algorithm is as follows:

- Use the message passing algorithm to max-calibrate a clique tree
- Find the most probable configuration 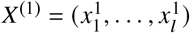
- For *i* = 2 → *M* − 1:
  1. Partition the whole configuration space into subspaces according to {*X*^(1)^, · · ·, *X*^(*i*−1)^} as described below
  2. Find the most probable configuration in each of the subspaces
  3. Compare all the maxima to find the next most probable configuration, *X*^(*i*+1)^

### Partitioning

*Given the ordering, define R*_*j*_ = *C*_*j*_*\S*_*j*_−_1_ *to be the variables left over by excluding the scope of sepset S*_*j*_−_1_ *from the scope of clique C*_*j*_ *(let R*_1_ = *C*_1_). For configuration *X*^*^ and *R*_*j*_, let the pair 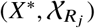 denote the subset of configuration space satisfying

a. *C*_*k*_ agrees with *X*^*^ for *k* = 1 to *j* − 1
b. *R*_*j*_ does not agree with *X*\*
c. *C*_*k*_, *k* ≥ *j* + 1 are unrestricted.

With this notation, the partitioning of the whole configuration space with respect to *X*^(1)^ is denoted as χ/*X*^(1)^ and can be expressed as

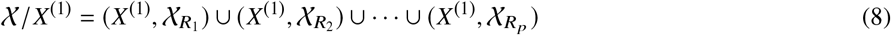

for *p* = |𝒞|. The partition with respect to {*X*^(1)^, *X*^(2)^} can then be created as follows. Note that *X*^(2)^ must belong to one of the subspaces in χ/*X*^(1)^; without loss of generality we assume it belongs to 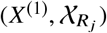. Then the partition of 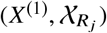 with respect to *X*^(2)^ is

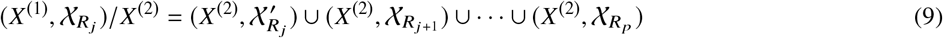

where 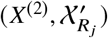 is the same subspace as 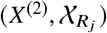, except *R*_*j*_ disagrees with both *X*^(1)^ and *X*^(2)^. The partition of the whole configuration space with respect to {*X*^(1)^, *X*^(2)^} is then the union of the sets (9) and (8) with 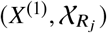 deleted:

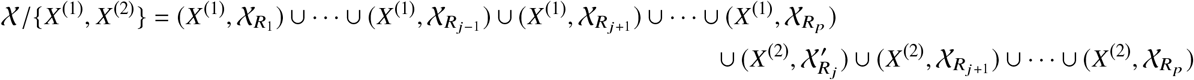

The partition with respect to {*X*^(1)^, · · ·, *X*^(*i*)^} is obtained similarly. Nilsson’s algorithm finds the maximum configuration over 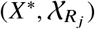, denoted *X*, as follows:

1. For *k* = 1 to *j* − 1, set *X*′ = *X*^*^ on *C*_*k*_
2. On *C*_*j*_, set *X* to

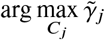

where 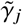is the potential γ_*j*_ with value set to zero on instantiations of *C*_*j*_ which agree with *X*\*
3. For the remaining cliques *k* ≥ *j* + 1, set *X* ′ on *C*_*k*_ to values derived using the max consistency property.

The maximum value of the unnormalized joint probability over 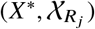 is also given by

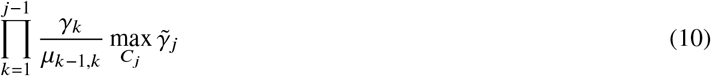

The enumeration algorithm proceeds by finding the configuration maximizing (10) among all pairs 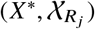 in the current partition, and then augmenting the partition according to the maximum. This can be implemented efficiently using a priority queue or heap data structure (34).

### Computational Complexity

Suppose the maximum number of configurations in any of the *p* cliques is *n*_*c*_. (For the helix-coil model presented here, *n*_*c*_ = 2^6^, and *p* = *l* − 5.) Then the maximum number of configurations in the sepsets and the residual set (*R*_*j*_) do not exceed *n*_*c*_. *Message passing:* Updating edge *i* → *j* requires *O*(*n*_*c*_) computations, so the total number of computations for calibration is *O*(*pn*_*c*_). *Finding maximum configurations:* Using the consistency property to find the maximum configuration in a pair 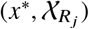 requires at most *n*_*c*_ + *pn*_*c*_ = *O*(*pn*_*c*_). Therefore finding the *M* most probable configurations requires *O*(*Mpn*_*c*_). *Partition calculations:* The number of new pairs 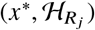 required to find the next probable configuration does not exceed *p*. Therefore the total number of pairs to find the *M* most probable configurations is *O*(*Mp*). For every pair, calculating the maximum value (10) requires at most *O*(*p* + *n*_*c*_) calculations. Inserting the *O*(*Mp*) pairs into a priority queue requires *O*(*Mp* log(*Mp*)) calculations. So the total number is *O*(*Mp*((*p* + *n*_*c*_) + log(*Mp*))). Altogether, the total number of calculations performed is

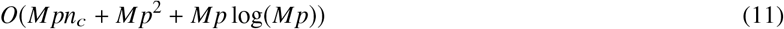

Therefore the complexity of Nilsson’s algorithm depends on the dominant term in (11). For the helix coil model, we have *n*_*c*_ = 2^6^ and *p* = *l−*5, so the total complexity of the algorithm is *O (Ml*^2^). In practice, the algorithm runs on peptides up to length 40 with α = 0.99 in approximately 100ms on an AMD A10-5745M APU.

## RESULTS

### Peptide Dataset

Helix-coil ensemble enumeration was performed for peptides from a large database of helical peptides constructed previously (18). The data set used here contains 1311 individual experimental conditions representing 361 distinct peptides; this represents approximately 95% of the data used to estimate the helix-coil model parameters, with a small number of peptides omitted from the analysis here due to length (6 ≤ or ≥ 40). Each helix-coil calculation takes as input the polypeptide sequence, a temperature, a solution pH, a solution urea concentration, a solution TMAO concentration, and indicators of N- and C-terminal blocking.

As this dataset is biased towards peptides with high helicity, a control dataset was constructed by substituting a randomly generated amino acid sequence for each peptide in the helical dataset (maintaining the experimental conditions intact). Random sequences were generated independently using amino acid frequencies obtained from the Expasy server (35). Figure 3 shows the distribution of the (model-predicted) mean helicities for this control dataset compared to the original helical peptide dataset. In what follows, we demonstrate several uses of ensemble enumeration to explore different aspects of helix coil models and modeling: (a) concentration of helix-coil populations, (b) the sources of error in previously proposed “single-sequence approximation” methods for calculating helicity, and (c) construction of visualization tools for exploring and comparing entire ensembles of peptides of interest. All of these applications are made possible by the HCEE algorithm’s ability to efficiently calculate highly accurate approximate ensembles.

**Figure 3.**
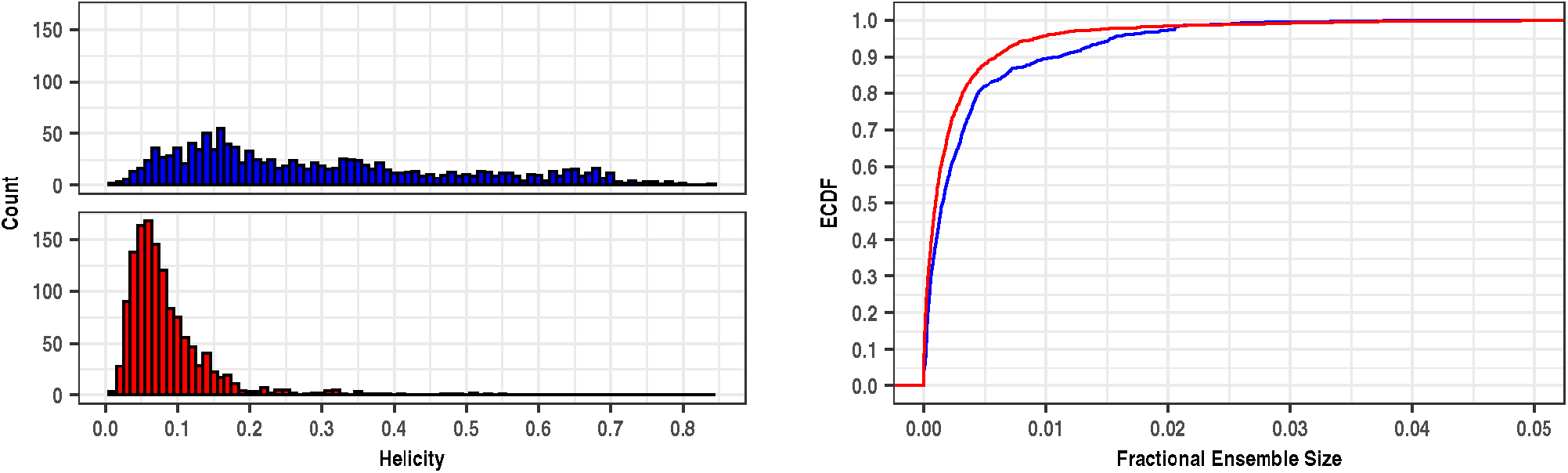
(a) Histogram showing the distribution of the predicted mean helicity for the helical peptide (blue) and control (red) datasets described in the text. The control peptides exhibit relatively uniform helicities, whereas the helical peptides have significantly higher mean helicity. (b) Empirical cumulative distribution function of the fractional ensemble size for the helical peptide (blue) and random peptide (red) datasets. For both datasets, we see that in over 95% of peptides, 99% of the peptide’s ensemble population can be accounted for by enumerating less than 2% of the total set of possible conformations.

**Figure 4.**
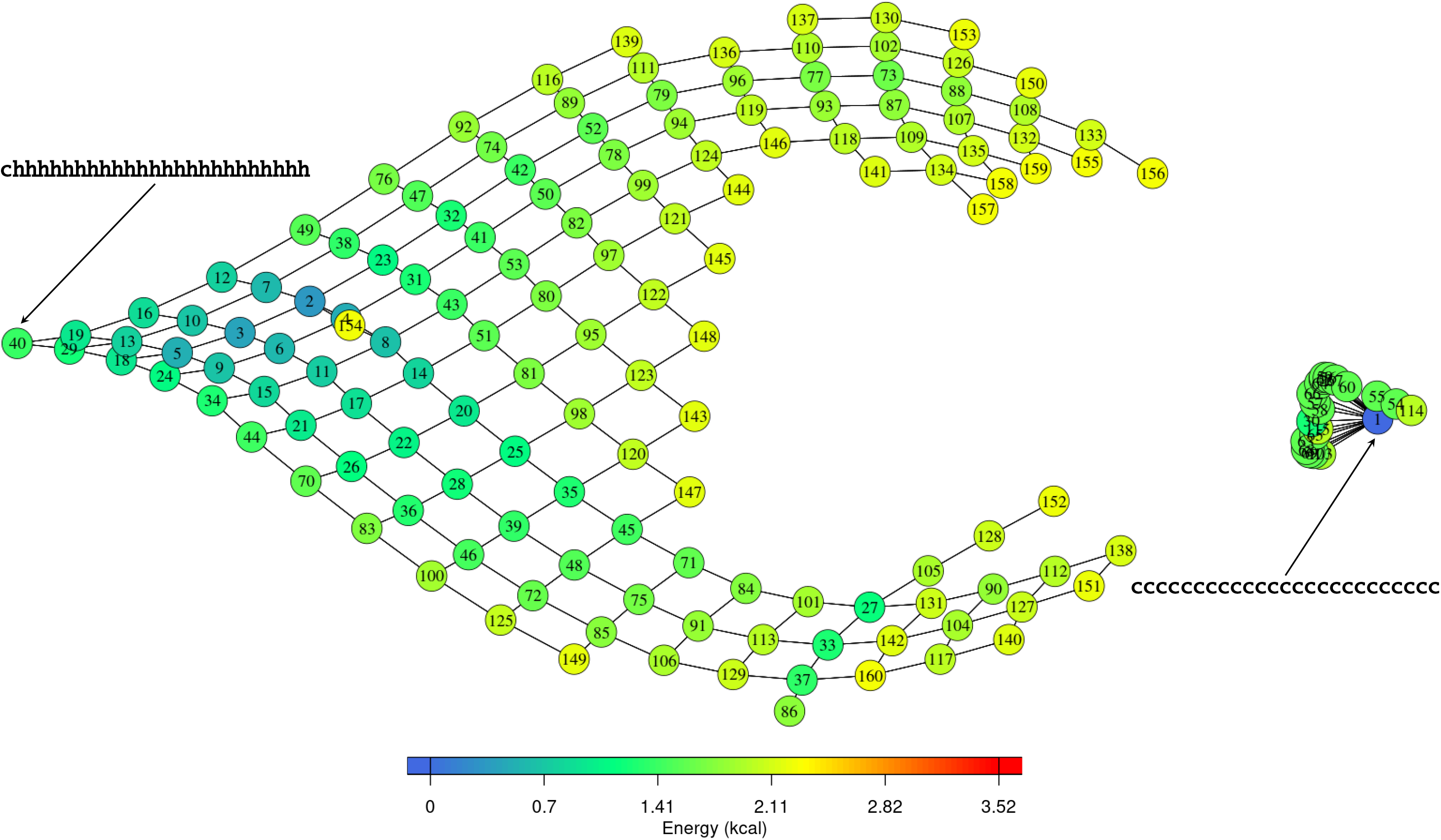
Plot of the graph for the alanine guest sequence from the Chan et. al (37) study. The vertices shown correspond to the minimal set of configurations that have a total aggregate population greater than 0.6. The vertices are labeled with the ranks of the associated configurations and colored according to the configuration free energy. The existence of an edge between two vertices means that the two configurations represented by the vertices differ by a Hamming distance of 1. The most helical and least helical configurations are labeled with the black arrows. The vertex partially occluded by the vertex labeled 154 is labeled 4. Here the multidimensional scaling has resulted in an arrangement of vertices that places more helical configurations towards the left and less helical configurations towards the right. The dense cluster of vertices around vertex 1 are configurations containing a single helical residue. The gap between this rightmost cluster and the rest of the graph is due to the high energy associated with having multiple helical residues that do not form stabilizing hydrogen bonds.

### Helix-coil Ensembles are Highly Concentrated

The HCEE (helix-coil ensemble enumeration) algorithm described in **Methods** was used to compute the Ψ_0.99_ approximate ensemble for each peptide input in both the helical and control datasets. Note that the value *α* = 0.99 ensures that all configurations lying within Ψ but outside Ψ_0.99_ collectively account for less than 1% of the population of the full ensemble.

Results from ensemble enumeration for the helical peptide dataset indicate that helix-coil ensembles place the vast majority of their probability on a small subset of the configuration space. Define the *fractional ensemble size f* Ψ_*α*_ as the proportion of the full configuration space occupied by Ψ_*α*_:

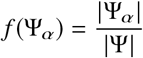

The fractional ensemble size provides a normalized measure of sub-ensemble size that can be used to compare the sizes of sub-ensembles derived from peptides of differing lengths. Figure 3 shows the empirical cdfs of fractional ensemble sizes obtained for both the helical peptide database and the random database.

We see that in both sets, the fraction of the available configuration space populated by the α = .99 sub-ensemble is relatively small. The mean fractional ensemble size seen for the combined datasets was 0.0032, and the 99% quantile was approximately 0.0278; that is, nearly 99% of the sub-ensembles populate less than 2.4% of the allowed configuration space. The maximum fractional ensemble size observed in either dataset was 0.145, obtained for the 8-residue random peptide ALILNDFR. Figure 3 also shows that the helical peptides exhibit larger fractional ensemble sizes on average than the random peptides. This may be explained in part by the high probability placed on the all-coil configuration within ensembles with low mean helicity.

This pattern of concentration on a relatively small subset persists within sub-ensembles (see Supplementary Materials). On average, the top 5% of configurations in a given Ψ_0.99_ sub-ensemble account for over 80% of the ensemble population, and the top 10% account for over 90% of the population. These results indicate a substantial degree of concentration *within* the Ψ_0.99_ sub-ensembles. (See Supplementary Materials for additional details.)

### Exploring Sources of Error in the Single-Sequence Approximation

The *single sequence approximation* is a simplification employed within AGADIR (10, 14–17) to reduce the computational complexity of helicity predictions, prior to the publication of an efficient algorithm for performing these calculations exactly(18). A *helical segment* is a sequence of contiguous residues with each residue taking on a helical (h) conformation. In the single sequence approximation, any configuration containing two or more non-overlapping helical segments is removed from the ensemble (i.e. each such configuration is assigned a statistical weight of zero) and the corresponding helicity calculation. This approximation is justified by arguing that such configurations must necessarily have very low probability. Additionally, AGADIR treats any helical segment of length three or fewer resdiues as coil; so only helical segments consisting of four or more helical residues contribute to the predicted helicity. Using the single sequence approximation, AGADIR then computes the helicity of a given peptide by explicitly enumerating all possible configurations containing a single helical segment of length four or more, and computing their statistical weights (14). The number of such configurations is substantially smaller than the full set of possible helix-coil configurations, dramatically reducing the computational cost of computing helicity under this approximation. A study to assess the accuracy of this single-sequence approximation by comparing to an alternative *multiple sequence* approximation (which includes configurations with multiple non-overlapping helical segments) found that the improvement in predictive accuracy from using the more expenseive multiple sequence approximation was negligible for peptides with a length shorter than 56 residues(17).

As exact calculation of the mean helicity over the full ensemble is now possible(18), exploring the sources of error in the single sequence approximation can provide some insight into the populations of helix-coil conformations. The HCEE algorithm developed here provides a way to do so: by enumerating the entire (1-*α* approximate) ensemble, we can identify the sources of error associated with ignoring configurations containing multiple non-overlapping helical segments.

To this end, the helix-coil model helicity prediction software was augmented to perform helicity calculations using the single sequence approximation alongside the full exact calculation. (Note that here, unlike AGADIR, we do not treat helical segments with a length less than four residues as coil segments; doing so will only increase the relative errors reported below.) Helicity predictions were then calculated using both exact and approximate approaches for all data points in the helical database, and the difference calculated to provide the error associated with the single sequence approximation for each data point. Some datapoints with high errors are shown in Table 1. Peptide L9 (PANLKALEAQKQKEQRQAAEELANAKKLKEQLEK) is a 34-residue fragment from Ribosomal protein L9 (36). Peptide EK (YSEEEEKKKKSSSEEEEKKKK) is one of a series of host-guest synthetic peptides with the host sequence `YSEEEEKKKK-XXX-EEEEKKKK` (28) and guest amino acid X. KDE (KDESYEELLRKTKAELLHWTKELTEEEKKALAEEGKIT) is a 38-residue peptide fragment of the Titin protein (14). Inspection of the Ψ_0.99_ sub-ensembles reveals that there are many high-ranking configurations containing multiple helical stretches. The most probable 100 configurations for the EK peptide and the KDE peptide are shown in Tables 4 and 5 respectively. The top 100 configurations of the EK peptide show a particularly striking pattern in which many of the configurations resemble a long helix broken in the middle around the area of the guest amino acid insertions. The top 100 configurations of the KDE peptide more often show configurations with a single helical segment, but there are still multiple high-ranking configurations that possess multiple helical segments of significant lengths. Enumeration of the ensembles for these peptides provides insight into the source of helicity prediction errors using the single sequence approximation, and shows that this approximation can significantly distort the view of the underlying helix-coil ensemble.

**Table 1:** Data points from helicity data base with the highest errors associated with the single sequence approximation.

| Peptide | Temp. | Exact Helicity | SS Approximation | Error |
| --- | --- | --- | --- | --- |
| L9 | 358 | 0.22 | 0.10 | -0.12 |
| KDE | 275 | 0.30 | 0.19 | -0.11 |
| EK | 277 | 0.36 | 0.25 | -0.11 |

**Table 2:**
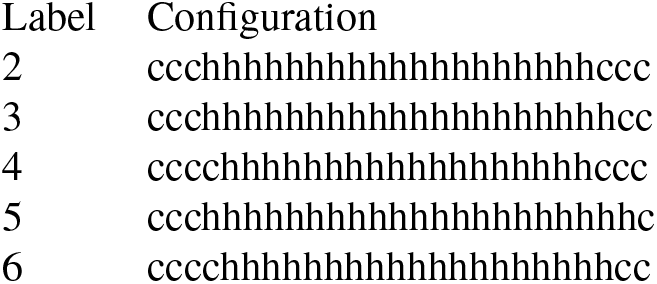
Configurations corresponding to the vertices within the basin centered around the local free energy minimum vertex labeled as 2 (the second highest populated configuration in the ensemble). All of these configurations contain coil in positions 2 and 3, with two of the five also containing coil at position 4. (Position 1 is restricted to coil due to absence of an N-terminal blocking group.)

**Table 3:**
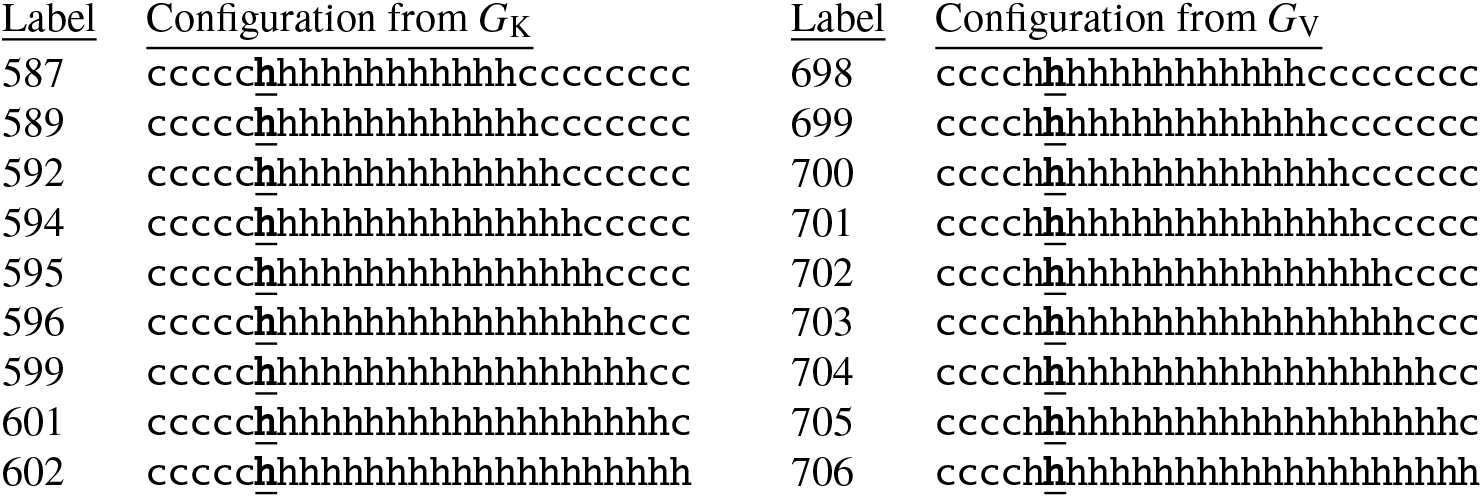
Configurations corresponding to the ‘double strip’ seen in the lower left portion of Figure 5. Position 6 (underlined) is the location of the first guest residue. The two parallel strips are seen to be mirror images of each other with only the configuration of residue 5 being different. These two configuration sequences represent two adjacent, alternate pathways between the all-helix and all-coil configurations as discussed in the text.

**Table 4:**
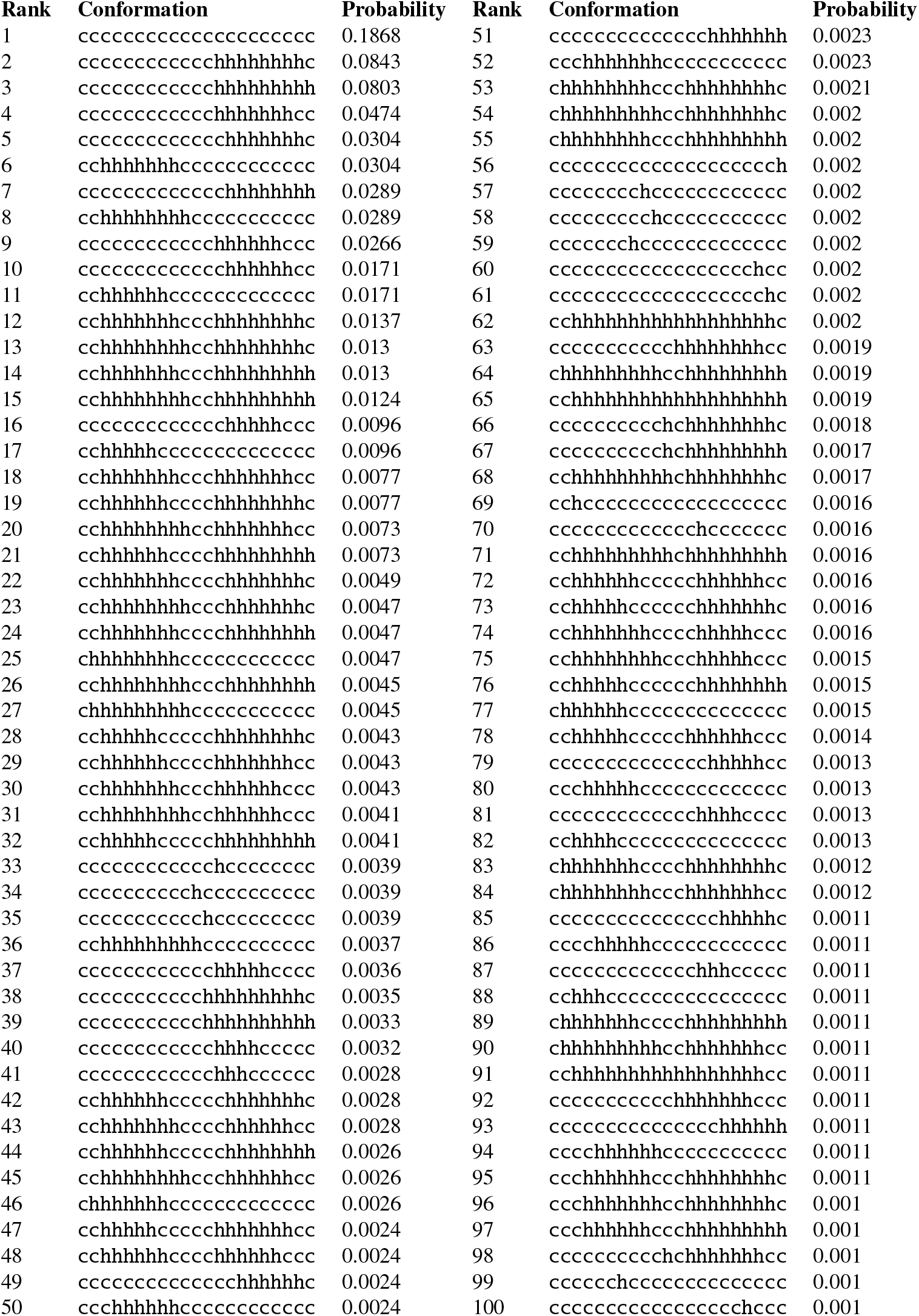
100 most probable configurations for the EK peptide. Probabilities have been rounded to four decimal places.

**Table 5:**
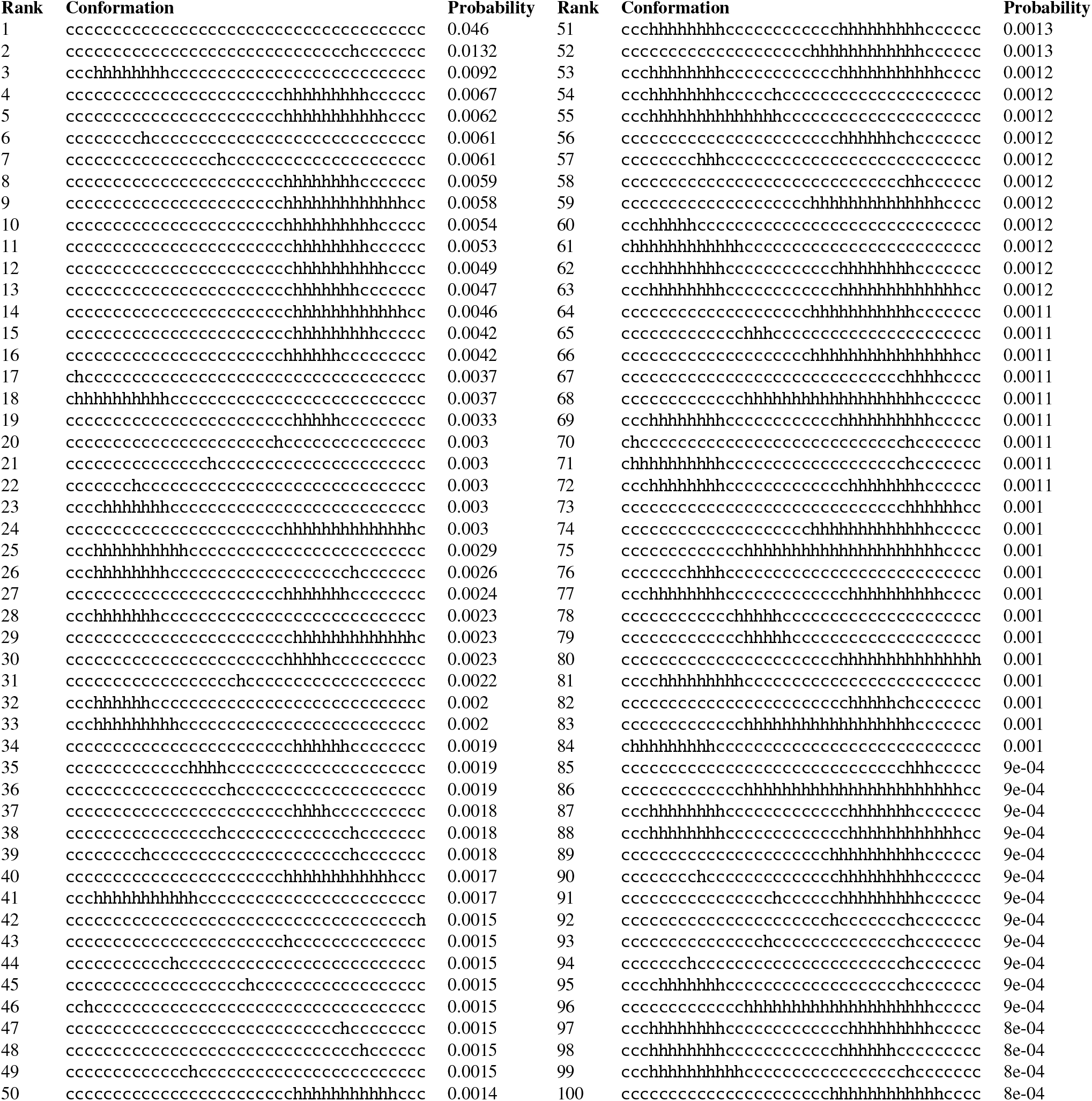
100 most probable configurations for the KDE peptide. Probabilities have been rounded to four decimal places.

### Visualizing Helix-coil Ensembles Using Graphs

The ability to enumerate an entire ((1 − *α*)-approximate) ensemble opens up interesting possibilities for visualization of helix-coil systems. Consider representing the configuration space of a helix-coil ensemble as a graph. Each possible configuration in the ensemble is represented as a vertex in the graph. An edge is placed between each pair of vertices whose configurations differ in exactly one position (Hamming distance equal to one). Recalling that there are at least 2^*n*−2^ such configurations for a peptide of length *n*, such a graph would be very difficult to visualize, even ignoring the exponential cost of computing it. Yet as shown above, a much smaller subset of the configuration space is actually significantly populated in a given ensemble, and therefore relevant to visualizing the system behavior. Enumerating the (1−α) -approximate ensemble Ψ_*α*_ which contains only the *m* most populated configurations of a given ensemble therefore provides a way of dramatically decreasing the complexity of the configuration space to be visualized, while retaining essentially all of the relevant information.

We consider the set of antimicrobial peptides studied by Chan et. al (37), who measured the average helicity of twenty designed 25-residue peptides at a temperature of 298 K and a pH of 7. Each peptide is an instance of the host sequence `KKAAAXAAAAAXAAWAAXAAAKKKK`, with ‘X’ denoting the guest residue. Each individual peptide has a single one of the twenty amino acids placed at the locations marked as ‘X’. All peptides have a C-terminal blocking group but no N-terminal blocking group, making `chhhhhhhhhhhhhhhhhhhhhhhh` the most helical configuration possible. The helix-coil model predicts the Chan helicity measurements well, with a mean squared error of 0.0059. (These measurements were included in the dataset used to fit the helix-coil model parameters.)

A minimal 0.9 sub-ensemble was generated for each peptide using the enumeration algorithm described in **Methods**; we denote the sub-ensemble for guest residue X by 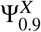. Computation of these sub-ensembles took only a few minutes on our hardware. The largest sub-ensemble was comprised of 179,648 configurations. Graphs were constructed from these sub-ensembles using the igraph library (38) of the R statistical package (39). The constructed graphs were visualized by projecting onto two dimensions using multidimensional scaling (40) (MDS) applied to the Hamming distance matrix.

### Visualizing the Alanine Guest Peptide Ensemble

Figure 4 shows the visualization obtained for the minimal ensemble 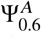 of the alanine guest peptide KKAAAAAAAAAAAAWAAAAAAKKKK. Each vertex is labeled with the ensemble rank of the configuration it represents, with the most probable configuration labeled as 1.Vertices are colored according to the configuration free energy under the helix-coil model, resulting in a visualization of the energy landscape over the ensemble of the alanine guest peptide (Figure 4).

The projection onto two dimensions preserves important topological structure of the ensemble. The x-axis is interpretable roughly as % helix, with the most helical configuration (configuration or rank 40) placed on the leftmost side of the plot and the all-coil configuration (conguration or rank 1) placed on the far right, and the percentage helicity decreasing as we move left to right in the graph. The effects of dimension reduction can be seen for example in the existence of vertices to the right of the all-coil configuration vertex, and the placement of configurations 4 and 154 nearly on top of each other. The dense cluster of vertices surrounding the all-coil configuration consists of (most of) the *n* − 1 configurations containing a single helical residue.

The graph is visibly separated into two connected components, indicating that paths between the primarily helical and primarily coil states must pass through configurations that are not highly populated; that is, they have free energies too high to be included in the ensemble 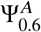.

The y-axis in the graph, on the other hand, appears to capture the N-to C-terminus dimension. For example, following paths originating at 40 proceeding towards 1 in the upper portion of the graph (such as those ending at 139, 150, and 156, whose configurations are shown in Table 6), we see that each of these paths is made up of a sequence of configurations containing a single helical stretch that is getting shorter from the C-terminal end. The paths differ in the number of coil residues N-terminal to the helix. Table 7 shows the configurations corresponding to similar paths in the lower half of the plot. We see the same phenomenon here, but now the isolated helical stretch is contracting from the N-terminus. Transitions from one path to an adjacent path correspond to adding or removing a helical residue at the opposite end of the helix from the contracting end.

**Table 6:**
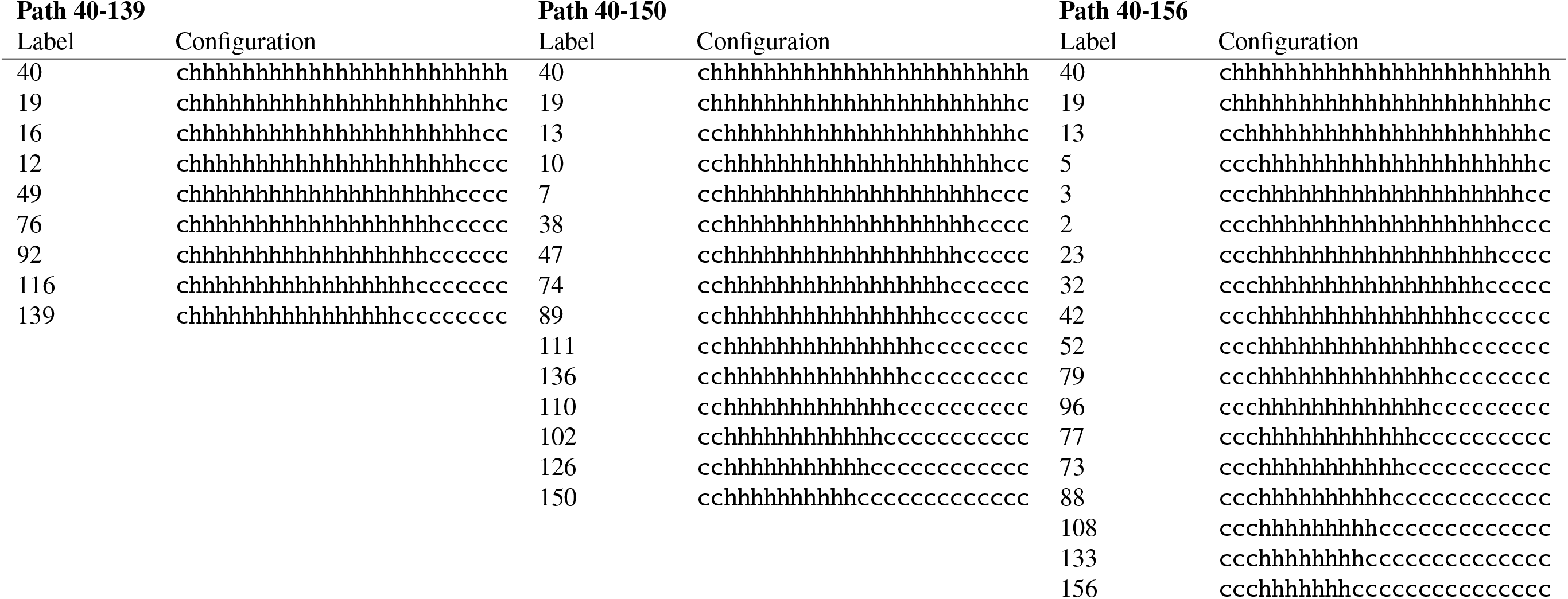
Configurations corresponding to three adjacent paths at the top of the graph in Figure 4, each beginning from the vertex labeled 40. Each path corresponds to the contraction of a single helix from the C-terminus to the N-terminus. Distinct paths correspond to distinct N-terminal helix starting positions.

**Table 7:**
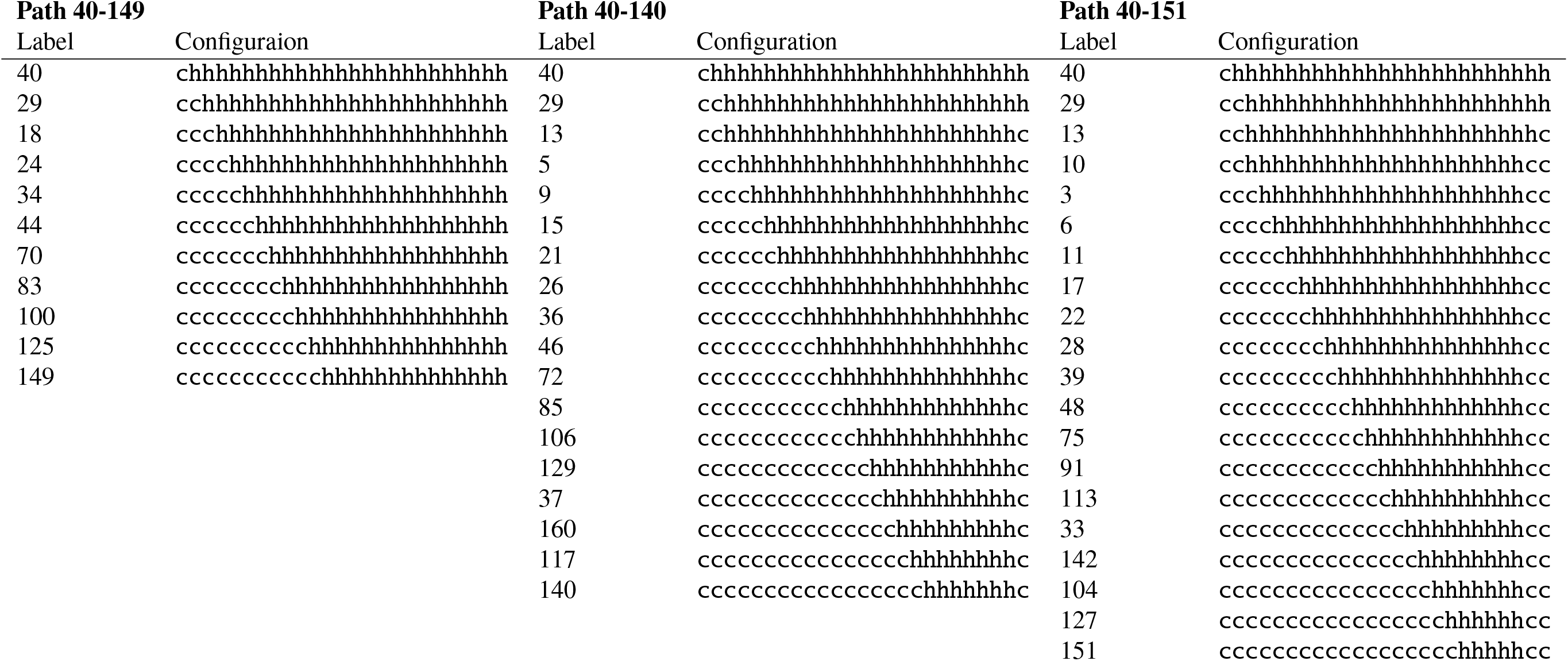
Configurations corresponding to three adjacent paths at the bottom of the graph in Figure 4. Each path corresponds to the contraction of a single helix from the N-terminus to the C-terminus. Distinct paths correspond to distinct C-terminal helix ending positions.

The coloring in Figure 4 also enables us to identify the presence of two local free energy minima in the ensemble, one at the all-coil configuration (vertex 1) - which is also the global free energy minimum - and the other appears to be located around vertex 2 on the more helical (left) side of the plot. Some of the vertices within this local minima are shown in Table 2. These configurations tend to have all residues helical except for the ends. This can be explained by energetic parameter values (not shown), where 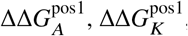, and ΔΔ*H*_*K*_ are all strongly positive while 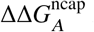 is strongly negative and 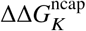 is zero. This strongly favors initiating the helix at an alanine preceded by another alanine (disfavoring starts at positions 2, 3, or 16, for example). 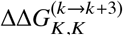 is also strongly positive, representing the unfavorable *k*→*k* + 3 interaction between the protonated sidechains of lysine, and the presence of this interaction between residues 22 and 25 combines with the positive ΔΔ*H*_K_ to disfavor the last four residues being helical.

### Visualizing the Effects of Perturbations Using Ensemble Difference Graphs

Graph-based ensemble visualizations can also be used to help study the effects that various perturbations may have on a helix-coil system. A useful tool for performing such an analysis is the *difference graph*. Let *G*_*i*_(*V*_*i*_, *E*_*i*_) and *G*_*k*_ (*V*_*k*_, *E*_*k*_) be two ensemble graphs. Define the difference graph Δ*G*_*ik*_ having vertex set

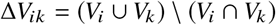

Here Δ*V*_*ik*_ contains only those vertices from *V*_*i*_ and *V*_*k*_ that do not appear in both *V*_*i*_ and *V*_*k*_. Similarly, the edge set for Δ*G*_*ik*_ is Δ*E*_*ik*_ and contains all edges in *E*_*i*_∪*E*_*k*_ having both endpoints in Δ*V*_*ik*_. If the underlying peptide sequences have the same length and terminal blocking states, then we can examine Δ*G*_*ik*_ to understand the effects of peturbation; for example, the ensembles represented by *G*_*i*_ and *G*_*k*_ may differ in their temperature, solvent pH, or sequence (e.g. one is a mutant of the other). The difference graph Δ*G*_*ik*_ then shows which configurations enter or leave the sub-ensemble (begin or cease to be significantly populated) of *G*_*i*_ (or *G*_*k*_) in response to the perturbation.

Ensemble graphs for the Chan study peptides were constructed at the α = 0.6 level, and difference graphs were generated for each pair of resulting graphs. Here *G*_*X*_ denotes the ensemble graph for the peptide with guest amino acid X, and Δ*G*_*XY*_ denotes the difference graph between *G*_*X*_ and *G*_*Y*_. Plots were generated for each ensemble graph and every difference graph. Each vertex was labeled with an integer that uniquely identifies its associated configuration in the union of the α = 0.6 ensembles.

Figure 5 shows the ensemble difference graph Δ*G*_KV_. Vertices found only in *G*_*K*_ are colored blue and vertices found only in *G*_*V*_ are colored green. Individual ensemble graphs *G*_K_ and *G*_V_ are also shown in Figure 5. A striking pattern observed in Δ*G*_KV_ is the existence of ‘double strips’ of contiguous connected vertices. These correspond to adjacent, alternate folding pathways included exclusively in one of the two subensemble graphs, indicating a shift in population from one pathway to the other. The configurations making up one of these double strips, spanning from vertices 698 and 587 to vertices 706 and 602, are shown in Table 3. We see that each vertex of the path from *G*_V_ has an isolated helical stretch that starts at residue 5, while each vertex of the path from *G*_K_ has an isolated helical stretch that starts at residue 6. Recall that residue 6 is the first location of the guest amino acid in the peptide sequence, so the configurations that appear in *G*_V_ have V in the second position of the helical stretch, and the configurations that appear in *G*_K_ have K in the first position of the helical stretch. Again, inspection of the energetic terms shows that the V → K perturbation significantly destabilizes the green vertex configurations in Figure 5 and stabilizes the blue vertex configurations. Similar explanations hold for the other ‘double-strips’ seen in Figure 5, i.e. (237-240,192-195), (46-48,34-36), and (544-577,676-692). This example demonstrates the utility of difference graph visualizations, enabled by the enumeration algorithm, to reveal deeper insights into the effects of perturbations on helix-coil ensembles.

**Figure 5.**
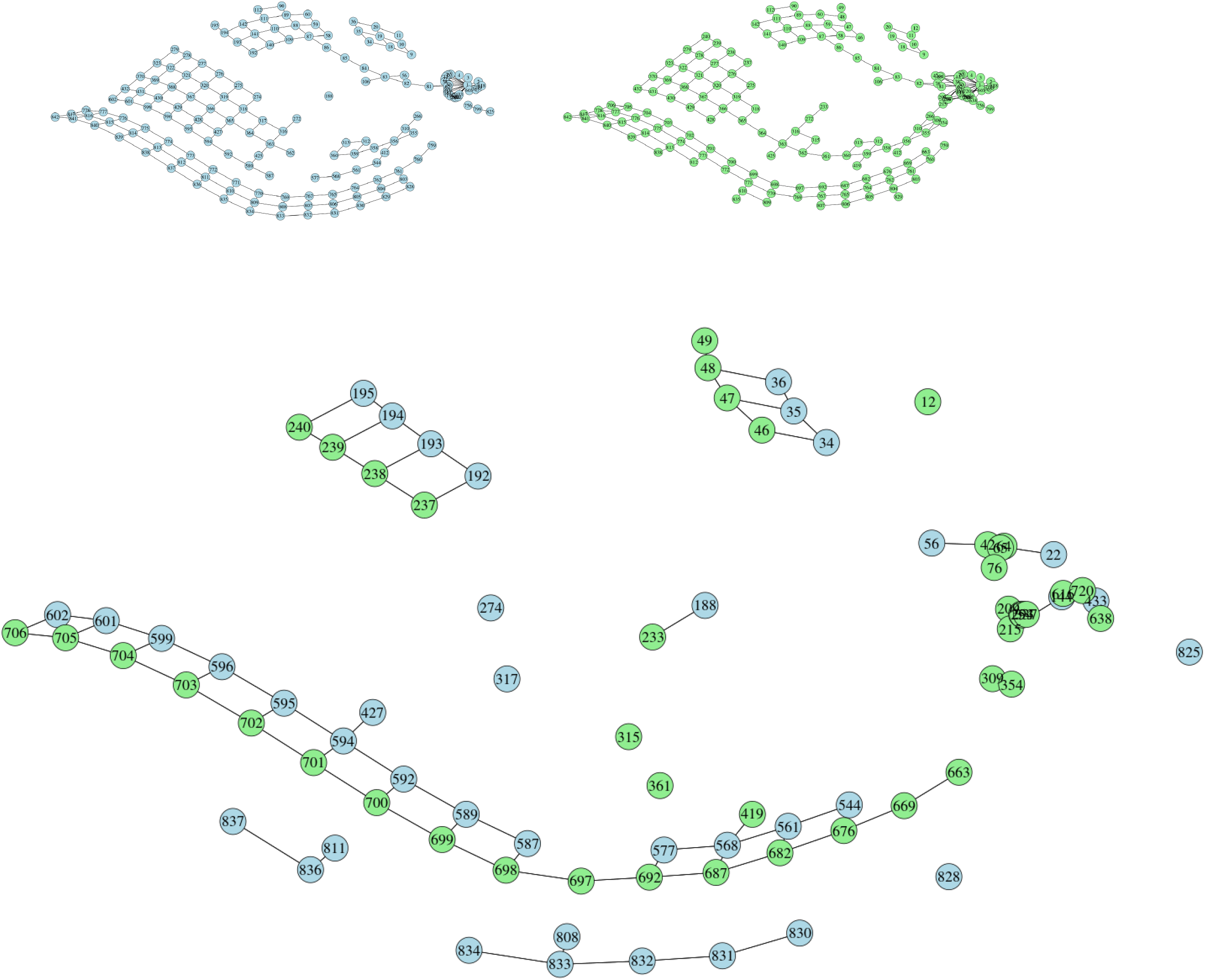
Plot of the difference graph (lower plot) Δ*G*_KV_, consisting of all vertices from *G*_K_ that do not appear in *G*_V_ (blue) and all vertices from *G*_V_ that do not appear in *G*_K_ (green). Plots of *G*_K_ and *G*_V_ are respectively shown in the upper left and upper right. Vertex labels uniquely identify each configuration in the union of the sub-ensembles generated for all the Chan study (37) peptides with *P*_total_ = 0.6. An edge between two vertices means that the two configurations differ by a Hamming distance of 1. Note the presence of the “double strips”, with each “strip” being of a different color.

### Insights on Helix Propagation Mechanisms from Helix-coil Ensemble Graphs

Analysis of the ensemble graphs produced using the enumeration algorithm has the potential to improve our current understanding of the dynamics of helix-coil ensembles. Here the dynamics of a helix-coil ensemble describes how a single peptide molecule in solution transitions between different configurations over time.

Visualization of ensemble graphs suggests other potential summaries of interest that may be computed from the enumerated (1−α) -approximate ensemble. For example, consider the problem of determining the most probable trajectory between the least helical and the most helical configurations in a helix-coil ensemble. We adopt a simple model whereby the energy of an edge (*i, j*) ∈ *E* is *E* ((*i, j*)) = *U*(*X* _*j*_) − *U*(*X*_*i*_), and the energy of a path λ is additive over the edges: *E*(λ) =Σ _(*i*_ ′_, *j*_′ _)∈*λ*_ *E*(*e*). This is consistent with e.g. a simple dynamics described by a weighted random-walk on the graph, where 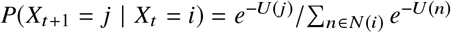, with the sum over the neighbors of *i* in the graph. The most probable (minimum energy) path between two vertices in the graph can then be found by applying a shortest path algorithm with edge weights

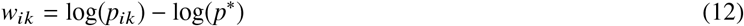

where *p*\* = min _(*r,s*) ∈ *E*_ log (*p*_*rs*_). In what follows the shortest path is obtained by Dijkstra’s algorithm using igraph (38). This most probable path represents a potential ‘folding pathway’ or mechanism for interconversion between all-helix and all-coil.

This interpretation bears some important caveats. Considerable previous experimental and theoretical work has provided a picture of helix-coil dynamics in partially helical peptides (41–44). In summary, helix nucleation is slow (100’s ns) relative to helix propagation (5 - 10 ns.) All of the edges of a helix-coil ensemble graph, except those from the rightmost all-coil vertex to any single-helix vertex, correspond to helix propagation. If we make the simplifying assumption that the rate of propagation is independent of position or sequence, then the graph proposes the sequence of steps that lead from the most populated single helix states to the all-helical state. By this assumption, the energy difference between adjacent vertices on an edge reflects the rate of the *h*→*c* transition (from left to right.) Taken together, these assumptions mean that a path created by edges between lowest energy configurations on a helix-coil ensemble graph represents a probable sequential mechanism for fluctuations between different configurations along the path. Then the most probable mechanism for interconversion between the all-helix and all-coil states is the shortest path on the graph.

Consider again the ensemble graph *G* _*A*_ for the alanine guest sequence from the Chan study (37) at approximation level α = 0.75. Recall that the least helical configuration for these peptides is ccccccccccccccccccccccccc and the most helical configuration is `chhhhhhhhhhhhhhhhhhhhhhhh`; denote the corresponding vertices in the ensemble graph by *v*_helix_ and *v*_coil_, respectively. Figure 6 depicts the most probable path for the alanine guest peptide from Chan et al. (37) Along this path, vertex 394 represents the highest energy configuration and the transitions to and from it are the likely rate-limiting transitions for the all-coil↔all-helix reaction. Thus, helix-formation in this sequence would include the rapid interconversion of all-coil and single h configurations followed by much slower nucleation of a helical turn at residues 19-21 that would rapidly propagate to configurations with 72-84% helicity. Of course this process is fully reversible, both macroscopically and microscopically.

**Figure 6.**
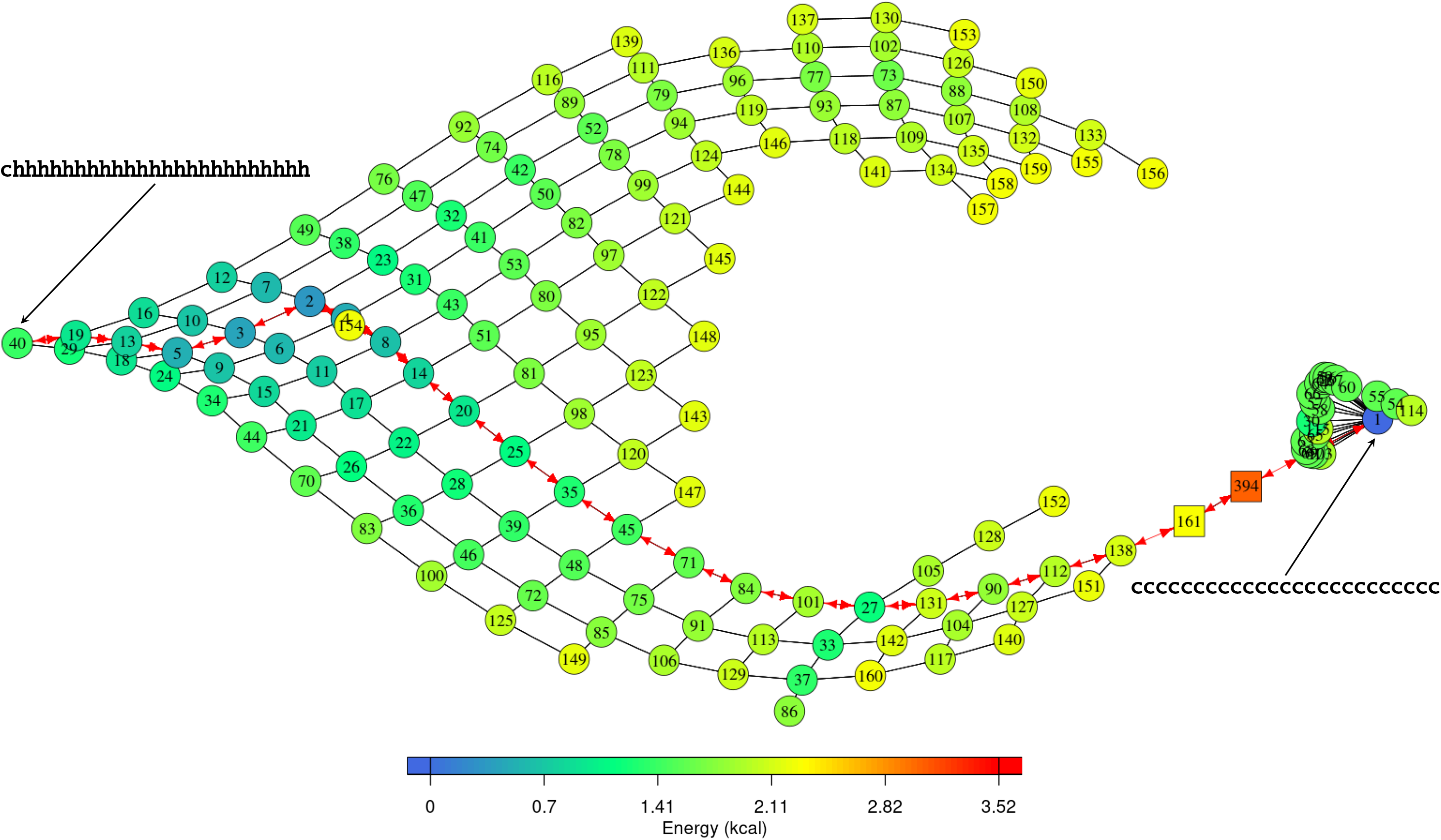
Plot showing the shortest path between the all-helix and all-coil configurations in the ensemble for the *P*_total_ = 0.75 ensemble graph of the alanine guest sequence from the Chan study (37). Circular vertices represent all configurations contained in the *P*_total_ = 0.6 ensemble graph. Square vertices show configurations lying outside the *P*_total_ = 0.6 ensemble but within the *P*_total_ = 0.75 ensemble that were necessary to form the shortest path. The shortest path (red double arrows) passes through the local free energy minima seen at vertices 2 and 27.

## DISCUSSION

The algorithm presented here enumerates configurations in order of decreasing probability, allowing characterization of the helix coil ensemble with high accuracy while avoiding the exponential exhaustive enumeration of the entire configuration space. In a large and representative collection of peptide sequences and conditions, we have found helix-coil ensembles to be highly concentrated, with the vast majority of probability mass being placed on a small subset of allowable configurations. This property allows approximation of the ensemble using a much smaller subset of configurations. This ensemble approximation approach allows for the effective enumeration of ensembles for purposes of investigating and visualizing collective properties of specific helix-coil ensembles.

An advantage of (approximate) ensemble enumeration over traditional helix-coil calculations is the provision of a convenient way to (approximately) calculate ensemble averages of collective properties beyond mean helicity (or other additive functionals). In addition to the applications shown here, helix-coil ensemble enumeration has several potential uses for further exploration. The last two decades have witnessed a rise in interest in the functional role of intrinsically disordered regions (IDR’s) in proteins.(45) The structural characterization of these regions presents many experimental and theoretical challenges. Because the formation of nascent helicity in these regions requires only local reduction in disorder, this type of intrachain interaction represents a highly probable transient structure in IDR’s. The use of the method presented here to predict the sequence dependence of nascent helicity and enumerate locations of the most likely helix represents a novel tool for understanding one feature of IDR structure.

Ensemble enumeration can also be useful in the design of artificial linkers. Intrinsically helical sequences have recently been used as so-called “rigid linkers.”(46) However, the sequences used for such linkers are usually repeats of EAAAK, which is not as helical as many authors assume. (For example, our estimate of the helicity of A(EAAAK)_3_A is only 36% at pH 7.5 and 300 K.) The methods described here provide useful insights into the distribution of helicity among the ensemble of configurations, which should aid in the rational design of linkers with specified rigidity.

## AUTHOR CONTRIBUTIONS

SZ and SS developed the enumeration algorithm. RH and SZ developed the software for performing calculations. TO advised on biophysical aspects including choosing examples and analyzing results. All authors contributed to writing the manuscript.

## ACKNOWLEDGEMENTS

This work was partially supported by NIH grant R01GM090201 (SCS) and NSF grant DMS1407622 (SCS) and the Duke University School of Medicine (TGO). The work was also supported by a Gates Millenium Scholars Graduate Fellowship provided through the Bill and Melinda Gates Foundation.

## REFERENCES

1. Poland, D., and H. A. Scheraga, 1970. Theory of Helix-Coil Transitions in Biopolymers. Academic Press.

2. Pauling, L., R. B. Corey, and H. R. Branson, 1951. The structure of proteins: Two hydrogen-bonded helical configurations of the polypeptide chain. Proc. Natl. Acad. Sci. U.S.A. 37:205–211.

3. Miller, W. G., D. A. Brant, and P. J. Flory, 1967. Random Coil Configurations of Polypeptide Copolymers. J. Mol. Biol. 23:67–80.

4. Zimm, B. H., and J. K. Bragg, 1959. Theory of the Phase Transition between Helix and Random Coil in Polypeptide Chains. J. Chem. Phys. 31:526–535.

5. Lifson, S., and A. Roig, 1961. On the Theory of Helix–Coil Transition in Polypeptides. J. Chem. Phys. 34:1963–1974.

6. Kent, S. B. H., 1988. Chemical Synthesis of Peptides and Proteins. Annu. Rev. Biochem. 57:957–989.

7. Scholtz, J. M., and R. L. Baldwin, 1992. The Mechanism of α-helix Formation by Peptides. Annu. Rev. Biophys. Biomol. Struct. 21:95–118.

8. Shalongo, W., and E. Stellwagen, 1995. Incorporation of pairwise interactions into the Lifson–Roig model for helix prediction. Protein Sci. 4:1161–1166.

9. Stapley, B. J., C. A. Rohl, and A. J. Doig, 1995. Addition of side chain interactions to modified Lifson–Roig helix–coil theory: Application to energetics of Phenylalanine–Methionine interactions. Protein Sci. 4:2383–2391.

10. Lacroix, E., A. R. Viguera, and L. Serrano, 1998. Elucidating the Folding Problem of α-helices: Local Motifs, Long-range Electrostatics, Ionic-strength Dependence and Prediction of NMR Parameters. J. Mol. Biol. 284:173–191.

11. Doig, A. J., A. Chakrabartty, T. M. Klingler, and R. L. Baldwin, 1994. Determination of Free Energies of N-Capping in α-Helices by Modification of the Lifson–Roig Helix–Coil Theory To Include N- and C-Capping. Biochemistry 33:3396–3403.

12. Park, S.-H., W. Shalongo, and E. Stellwagen, 1993. Residue Helix Parameters Obtained from Dichroic Analysis of Peptides of Defined Sequence. Biochemistry 32:7048–7053.

13. Park, S.-H., W. Shalongo, and E. Stellwagen, 1993. Modulation of the Helical Stability of a Model Peptide by Ionic Residues. Biochemistry 32:12901–12905.

14. Muñoz, V., and L. Serrano, 1995. Elucidating the Folding Problem of Helical Peptides using Empirical Parameters. II. Helix Macrodipole Effects and Rational Modification of the Helical Content of Natural Peptides. J. Mol. Biol. 245:275–296.

15. Muñoz, V., and L. Serrano, 1995. Elucidating the Folding Problem of Helical Peptides using Empirical Parameters. III. Temperature and pH Dependence. J. Mol. Biol. 245:297–308.

16. Muñoz, V., and L. Serrano, 1994. Elucidating the folding problem of helical peptides using empirical parameters. Nat. Struct. Mol. Biol. 1:399–409.

17. Muñoz, V., and L. Serrano, 1997. Development of the Multiple Sequence Approximation Within the AGADIR Model of α-helix Formation: Comparison with Zimm–Bragg and Lifson–Roig Formalisms. Biopolymers 41:495–509.

18. Schmidler, S. C., J. E. Lucas, and T. G. Oas, 2007. Statistical Estimation of Statistical Mechanical Models: Helix-Coil Theory and Peptide Helicity Prediction. J. Comput. Biol. 14:1287–1310.

19. Raucci, R., G. Colonna, G. Castello, and S. Costantini, 2013. Peptide Folding Problem: A Molecular Dynamics Study on Polyalanines Using Different Force Fields. Int. J. Pept. Res. Ther. 19:117–123.

20. Cooke, B., and S. C. Schmidler, 2008. Statistical Prediction and Molecular Dynamics Simulation. Biophys. J. 95:4497–4511.

21. Best, R. B., and G. Hummer, 2009. Optimized molecular dynamics force fields applied to the helix-coil transition of polypeptides. J. Phys. Chem. B 113:9004–9015.

22. Chen, Y.-H., J. T. Yang, and K. H. Chau, 1974. Determination of the Helix and β Form of Proteins in Aqueous Solution by Circular Dichroism. Biochemistry 13:3350–3359.

23. Jerrum, M., and A. Sinclair, 1993. Polynomial-Time Approximation Algorithms for the Ising Model. SIAM J. Comput. 22:1087–1116.

24. Georgiev, I., R. H. Liliean, and B. Donald, 2008. The Minimized Dead-End Elimination Criterion and Its Application to Protein Redesign in a Hybrid Scoring and Search Algorithm For Computing Partition Functions Over Molecular Ensembles. J. Comp. Chem. 29:1527–42.

25. Huyghues-Despointes, B. M. P., T. M. Klingler, and R. L. Baldwin, 1995. Measuring the Strength of Side-Chain Hydrogen Bonds in Peptide Helices : The Gln · Asp (i, i + 4) Interaction. Biochemistry 34:13267–13271.

26. Marqusee, S., and R. L. Baldwin, 1987. Helix stabilization by Glu^−^ Lys^+^ salt bridges in short peptides of de novo design. Proc. Natl Acad. Sci. U.S.A. 84:8898–8902.

27. Padmanabhan, S., and R. L. Baldwin, 1994. Tests for helix-stabilizing interactions between various nonpolar side chains in alanine-based peptides. Protein Sci. 3:1992–1997.

28. Lyu, P. C., M. I. Liff, L. A. Marky, and N. R. Kallenbach, 1990. Side Chain Contributions to the Stability of Alpha-Helical Structure in Peptides. Science 250:669–673.

29. Lauritzen, S. L., 1996. Graphical Models. Oxford.

30. Koller, D., and N. Friedman, 2009. Probabilistic Graphical Models: Principles and Techniques. The MIT Press, 1 edition.

31. Bilmes, J. A., and C. Bartels, 2005. Graphical Model Architectures for Speech Recognition. IEEE Signal Process. Mag. 22:89–100.

32. Sachs, K., O. Perez, D. Pe’er, D. A. Lauffenburger, and G. P. Nolan, 2005. Causal Protein-Signaling Networks Derived from Multiparameter Single-Cell Data. Sci. Signal 308:523.

33. Nilsson, D., 1998. An efficient algorithm for finding the M most probable configurations in probabilistic expert systems. Stat. Comput. 8:159–173.

34. Cormen, T. H., C. E. Leiserson, R. L. Rivest, and C. Stein, 2009. Introduction to Algorithms, Third Edition. The MIT Press, 3rd edition.

35. A., B. Release notes for UniProtKB/Swiss-Prot release 2013_04 - April 2013. https://web.expasy.org/protscale/pscale/A.A.Swiss-Prot.html.

36. Kuhlman, B., H. Y. Yang, J. A. Boice, R. Fairman, and D. P. Raleigh, 1997. An Exceptionally Stable Helix from the Ribosomal Protein L9: Implications for Protein Folding and Stability. J. Mol. Biol. 270:640–647.

37. Chan, C., L. L. Burrows, and C. M. Deber, 2004. Helix Induction in Antimicrobial Peptides by Alginate in Biofilms. J. Biol. Chem. 279:38749–38754.

38. Csárdi, G., and T. Nepusz, 2006. The igraph software package for complex network research. InterJournal Complex Systems:1695. https://igraph.org.

39. R Core Team, 2023. R: A Language and Environment for Statistical Computing. R Foundation for Statistical Computing, Vienna, Austria. https://www.R-project.org/.

40. Cox, T. F., and M. A. A. Cox, 2001. Multidimensional Scaling. Chapman and Hall, 2^nd^ edition.

41. Hammes, G. G., and P. B. Roberts, 1969. Dynamics of the Helix–coil Transition in Poly-L-ornithine. J. Am. Chem. Soc. 91:1812–1816.

42. Thompson, P. A., V. Muñoz, G. S. Jas, E. R. Henry, W. A. Eaton, and J. Hofrichter, 2000. The Helix-Coil Kinetics of a Heteropeptide. J. Phys. Chem. B 104:378–389.

43. Wang, T., Y. Zhu, Z. Getahun, D. Du, C.-Y. Huang, W. F. DeGrado, and F. Gai, 2004. Length Dependent Helix–Coil Transition Kinetics of Nine Alanine-Based Peptides. J. Phys. Chem. B 108:15301–15310.

44. Poland, D., and H. A. Scheraga, 1966. Kinetics of the Helix–Coil Transition in Polyamino Acids. J. Chem. Phys. 45:2071–2090.

45. Uversky, V. N., 2015. Functional roles of transiently and intrinsically disordered regions within proteins. FEBS J. 282:1182–1189.

46. Arai, R., H. Ueda, A. Kitayama, N. Kamiya, and T. Nagamune, 2001. Design of the linkers which effectively separate domains of a bifunctional fusion protein. Protein Engineering, Design and Selection 14:529–532.

