## Supplementary Material for "Efficient Enumeration and Visualization of Helix-Coil Ensembles"

### Helix-coil model

The energy of a configuration $X$, relative to the reference (all-coil) configuration, is given by

$$U(X,\Omega) = \sum_{i=1}^{l} \Delta G_i(X,\Omega) + \sum_{i=1}^{l}\sum_{k=1}^{3} \Delta\Delta G_{i-k:i+k}(X,\Omega) + \Delta\Delta G_{\mathrm{term}}(X,\Omega)$$

where $\Delta G_i$ is the contribution of residue $i$ to the free energy relative to the reference state, and $\Delta\Delta G_{i-k,i+k}$ contains additional energetic contributions from sidechain-sidechain and sidechain-backbone interactions spanning the corresponding window of residues (of length $2k+1$) centered at sequence position $i$. More specifically, we have:

$$\Delta G_i(X,\Omega) = \Delta H_{\omega_i} x_{i-1:i+1} - T\Delta S_{\omega_i} x_i + [\mathrm{Urea}]m_{\mathrm{u}} \cdot x_{i-1:i+1} + [\mathrm{Tmao}]m_{\mathrm{t}} \cdot x_{i-1:i+1}$$

where $\bar{x} = 1 - x$ and $x_{j:k} = \prod_{i=j}^{k} x_i$. (We define $x_i = 0$ for any $i < 1$ or $i > l$.) Here

$$\Delta H_{\omega_i} = \Delta H + \Delta\Delta H_{\omega_i} + \Delta C_{p,\omega_i}(T - T_0) \qquad \Delta S_{\omega_i} = \Delta S^{n}_{\omega_i} + (1 - f^{\mathrm{n}}_{\omega_i})\Delta\Delta S_{\omega_i} + \Delta C_{p,\omega_i} \log\left(\frac{T}{T_0}\right)$$

are the intrinsic enthalpy and entropy of helix formation for residue $i$, and

$$\Delta\Delta H_{\omega_i} = f^{+}_{\omega_i}\Delta\Delta H^{+}_{\omega_i} + (1 - f^{+}_{\omega_i})\Delta\Delta H^{-}_{\omega_i}$$

with $f^{+}_{\omega_i} = (1 + 10^{\mathrm{pH}-\mathrm{pKa}_i})^{-1}$ being the fractional population of the protonated form of the sidechain for residue $i$, and $f^{\mathrm{n}}_{\omega_i}$ the fractional population of the sidechain protonation state of residue $i$ which is most abundant at neutral pH. $\Delta S^{n}_{\omega_i}$ is the intrinsic change in entropy experienced by a residue when it goes from coil to helix and the sidechain protonation state is consistent with that most abundant at neutral pH. $\Delta\Delta S_{\omega_i} = \Delta S^{\bar{n}}_{\omega_i} - \Delta S^{n}_{\omega_i}$, where $\Delta S^{\bar{n}}_{\omega_i}$ is the intrinsic change in entropy experienced by a residue when it goes from coil to helix and the sidechain protonation state is consistent with the one that is less abundant at neutral pH.

The heat capacity change experienced by a residue when it goes from coil to helix is given by:

$$\Delta C_{p,\omega_i} = \Delta C_p + f^{+}_{\omega_i}\Delta\Delta C^{+}_{p,\omega_i} + (1 - f^{+}_{\omega_i})\Delta\Delta C^{-}_{p,\omega_i}$$

at temperature $T$, with reference temperature $T_0 = 273$K. This temperature dependence of $\Delta H_{\omega_i}$ and $\Delta S_{\omega_i}$ is largely inspired by the temperature dependence defined within AGADIR (1). $m_{\mathrm{u}}$ and $m_{\mathrm{t}}$ account for the sidechain-independent effects of the osmolytes urea and TMAO, respectively, on the residue-specific change in free energy upon helix formation (2, 3).

The $\Delta\Delta G$ interaction terms consist of the following:

$$\begin{aligned}
\Delta\Delta G_{i-1:i+1}(X,\Omega) &= \Delta\Delta G^{\mathrm{Ncap}}_{\omega_i} \cdot \bar{x}_i x_{i+1} + \Delta\Delta G^{\mathrm{pos1}}_{\omega_i} \cdot \bar{x}_{i-1} x_i + \Delta\Delta G^{\mathrm{CP}}_{\omega_i,\omega_{i+1}} \cdot x_{i-1}\bar{x}_{i:i+1} \\
\Delta\Delta G_{i-2:i+2}(X,\Omega) &= \Delta\Delta G^{\mathrm{pos2}}_{\omega_i} \cdot \bar{x}_{i-2} x_{i-1:i} + \Delta\Delta G^{\mathrm{ii3}}_{\omega_{i-2},\omega_{i+1}} \cdot x_{i-2:i+1} + \Delta\Delta G^{\mathrm{ii4}}_{\omega_{i-2},\omega_{i+2}} \cdot x_{i-2:i+2} + \Delta\Delta G^{\mathrm{CB}}_{\omega_{i-2},\omega_{i+1}} \cdot \bar{x}_{i-2} x_{i-1:i+1} \\
&\quad + \Delta\Delta G^{\mathrm{CA1}}_{\omega_{i-2},\omega_{i+1}} \cdot \bar{x}_{i-2} x_{i-1:i+1} + \Delta\Delta G^{\mathrm{CS}}_{\omega_{i-2},\omega_{i+2},\omega_{i+1}} \cdot x_{i-2:i}\bar{x}_{i+1:i+2} + \Delta\Delta G^{\mathrm{CA3a}}_{\omega_{i-2},\omega_{i+2}} \cdot x_{i-2:i+1}\bar{x}_{i+2} \\
\Delta\Delta G_{i-3:i+3}(X,\Omega) &= \Delta\Delta G^{\mathrm{CH}}_{\omega_{i-3},\omega_{i+2},\omega_{i-2}} \cdot \bar{x}_{i-3:i-2} x_{i-1:i+2} + \Delta\Delta G^{\mathrm{CA2}}_{\omega_{i-2},\omega_{i+2}} \cdot \bar{x}_{i-3:i-2} x_{i-1:i+2} + \Delta\Delta G^{\mathrm{CA3b}}_{\omega_{i-3},\omega_{i+1}} \cdot x_{i-3:i+1}\bar{x}_{i+2} \\
\Delta\Delta G_{\mathrm{term}}(X,\Omega) &= \Delta\Delta G^{\mathrm{Nterm}} \cdot b_N x_1 + \Delta\Delta G^{\mathrm{CBN}}_{\omega_1,\omega_4,\omega_5} \cdot \bar{b}_N \bar{x}_1 x_{2:5}
\end{aligned}$$

The individual parameters comprising these interaction terms are described in Table 8. Terms involving ionizable sidechains are split into two contributions based on the side chain protonation state, e.g.

$$\Delta\Delta G^{\mathrm{Ncap}}_{\omega_i} = f^{+}_{\omega_i}\Delta\Delta G^{\mathrm{Ncap+}}_{\omega_i} + (1 - f^{+}_{\omega_i})\Delta\Delta G^{\mathrm{Ncap\text{-}}}_{\omega_i}$$

Parameters $\Delta\Delta G^{\mathrm{ii3}}_{A,B}$ represent energies associated with $i \longrightarrow i+3$ sidechain-sidechain interactions between residues of amino acid types $A$ and $B$ (separated by $3 - 1 = 2$ intervening residues). These interactions are represented by (up to) three distinct parameters, depending on whether both sidechains are protonated ($\Delta\Delta G^{\mathrm{ii3++}}_{A,B}$), only one protonated ($\Delta\Delta G^{\mathrm{ii3+-}}_{A,B}$), or both deprotonated ($\Delta\Delta G^{\mathrm{ii3--}}_{A,B}$). A sidechain that is not pH-dependent is always considered to be deprotonated. The parameters $\Delta\Delta G^{\mathrm{ii4}}_{A,B}$ are defined similarly for $i \longrightarrow i+4$ interactions.

| **Parameter** | **Free energy contribution of:** |
|---|---|
| $\Delta\Delta G^{\text{Ncap}}_{\omega_j}$ | Amino acid residue of type $\omega_j$ in coil state immediately preceding start of a helix (4–6). |
| $\Delta\Delta G^{\text{pos1}}_{\omega_j}$ | Amino acid residue of type $\omega_j$ in the first position of a helix (7–9). |
| $\Delta\Delta G^{\text{pos2}}_{\omega_j}$ | Amino acid residue of type $\omega_j$ in the second position of a helix (7–9). |
| $\Delta\Delta G^{\text{Nterm}}$ | Helix stabilizing effect of acetylating (blocking) the peptide N-terminus on the interaction of the first residue with the helix dipole (10). |
| $\Delta\Delta G^{\text{ii3}}_{\omega_{j-2},\omega_{j+1}}$ | Long range sidechain-sidechain interaction between residues of types $\omega_{j-2}$ and $\omega_{j+1}$ occurring when residues $j-2$ through $j+1$ are all helical (11–16, 16, 17). |
| $\Delta\Delta G^{\text{ii4}}_{\omega_{j-2},\omega_{j+2}}$ | Long range sidechain-sidechain interaction between residues of types $\omega_{j-2}$ and $\omega_{j+2}$ occurring when residues $j-2$ through $j+2$ are all helical (11–16, 16, 17). |
| $\Delta\Delta G^{\text{CB}}_{\omega_{j-2},\omega_{j+1}}$ | Capping box motif with residues of types $\omega_{j-2}$ and $\omega_{j+1}$ in the Ncap and N3 positions respectively (18). |
| $\Delta\Delta G^{\text{CBN}}_{\omega_1,\omega_4,\omega_5}$ | Additional contribution to the capping box when it appears at the unblocked N-terminus of the peptide (19). |
| $\Delta\Delta G^{\text{CH}}_{\omega_{j-3},\omega_{j+2},\omega_{j-2}}$ | Hydrophobic staple motif with residues of types $\omega_{j-3}$,$\omega_{j+2}$, and $\omega_{j-2}$ in the N', N4, and Ncap positions respectively (20, 21). |
| $\Delta\Delta G^{\text{CS}}_{\omega_{j-2},\omega_{j+2},\omega_{j+1}}$ | Schellman motif with residues of types $\omega_{j-2}$, $\omega_{j+2}$ , and $\omega_{j+1}$ in the C3, C', and Ccap positions respectively (22–24). |
| $\Delta\Delta G^{\text{CP}}_{\omega_j,\omega_{j+1}}$ | Pro C-capping motif with residues of types $\omega_j$ and $\omega_{j+1}$ in the Ccap and C' positions respectively (25). |
| $\Delta\Delta G^{\text{CA1}}_{\omega_{j-2},\omega_{j+1}}$ | Capping box motif variant described in AGADIR (26) with residues of types $\omega_{j-2}$ and $\omega_{j+1}$ in positions Ncap and N3 respectively. |
| $\Delta\Delta G^{\text{CA2}}_{\omega_{j-2},\omega_{j+2}}$ | Capping motif described in AGADIR (26), similar to the hydrophobic staple motif, with residues of types $\omega_{j-2}$ and $\omega_{j+2}$ in positions N4 and Ncap respectively. |
| $\Delta\Delta G^{\text{CA3a}}_{\omega_{j-2},\omega_{j+2}}$ | Capping motif described in AGADIR (26) involving an aromatic interaction, with residues of types $\omega_{j-2}$ and $\omega_{j+2}$ in positions C4 and Ccap respectively. |
| $\Delta\Delta G^{\text{CA3b}}_{\omega_{j-3},\omega_{j+1}}$ | Capping motif described in AGADIR (26) involving an aromatic interaction, with residues of types $\omega_{j-3}$ and $\omega_{j+1}$ in positions C5 and C1 respectively. |

Table 8: Descriptions of helix coil model parameters that appear in the $\Delta\Delta G_{i-k:i+k}$ and $\Delta\Delta G_{\text{term}}$ terms of the energy function (Equation 1).

### SUPPLEMENTARY MATERIAL REFERENCES

[S1] Muñoz, V., and L. Serrano, 1995. Elucidating the Folding Problem of Helical Peptides using Empirical Parameters. III. Temperature and pH Dependence. J. Mol. Biol. 245:297–308.

[S2] Scholtz, J. M., D. Barrick, E. J. York, J. M. Stewart, and R. L. Baldwin, 1995. Urea unfolding of peptide helices as a model for interpreting protein unfolding. Proc. Natl. Acad. Sci. U.S.A. 92:185–189.

[S3] Celinski, S. A., and J. M. Scholtz, 2002. Osmolyte effects on helix formation in peptides and the stability of coiled-coils. Protein Sci. 11:2048–2051.

[S4] Doig, A. J., A. Chakrabartty, T. M. Klingler, and R. L. Baldwin, 1994. Determination of Free Energies of N-Capping in $\alpha$-Helices by Modification of the Lifson–Roig Helix–Coil Theory To Include N- and C-Capping. Biochemistry 33:3396–3403.

[S5] Doig, A. J., and R. L. Baldwin, 1995. N- and C-capping preferences for all 20 amino acids in $\alpha$-helical peptides. Protein Sci. 4:1325–1336.

[S6] Aurora, R., and G. D. Rose, 1998. Helix capping. Protein Sci. 7:21–38.

[S7] Richardson, J. S., and D. C. Richardson, 1988. Amino Acid Preferences for Specific Locations at the Ends of $\alpha$ Helices. Science 240:1648–1652.

[S8] Presta, L. G., and G. D. Rose, 1988. Helix Signals in Proteins. Science 240:1632–1641.

[S9] Schmidler, S. C., J. S. Liu, and D. L. Brutlag, 2000. Bayesian Segmentation of Protein Secondary Structure. J. Comput. Biol. 7:233–248.

[S10] Marqusee, S., and R. L. Baldwin, 1987. Helix stabilization by $Glu^- \cdots Lys^+$ salt bridges in short peptides of *de novo* design. Proc. Natl Acad. Sci. U.S.A. 84:8898–8902.

[S11] Padmanabhan, S., and R. L. Baldwin, 1994. Tests for helix-stabilizing interactions between various nonpolar side chains in alanine-based peptides. Protein Sci. 3:1992–1997.

[S12] Klingler, T. M., and D. L. Brutlag, 1994. Discovering structural correlations in $\alpha$-helices. Protein Sci. 3:1847–1857.

[S13] Creamer, T. P., and G. D. Rose, 1995. Interactions between hydrophobic side chains within $\alpha$-helices. Protein Sci. 4:1305–1314.

[S14] Huyghues-Despointes, B. M. P., T. M. Klingler, and R. L. Baldwin, 1995. Measuring the Strength of Side-Chain Hydrogen Bonds in Peptide Helices : The Gln · Asp $(i, i + 4)$ Interaction. Biochemistry 34:13267–13271.

[S15] Shalongo, W., and E. Stellwagen, 1995. Incorporation of pairwise interactions into the Lifson–Roig model for helix prediction. Protein Sci. 4:1161–1166.

[S16] Stapley, B. J., C. A. Rohl, and A. J. Doig, 1995. Addition of side chain interactions to modified Lifson–Roig helix–coil theory: Application to energetics of Phenylalanine–Methionine interactions. Protein Sci. 4:2383–2391.

[S17] Fernández-Recio, J., A. Vázquez, C. Civera, P. Sevilla, and J. Sancho, 1997. The Tryptophan/Histidine Interaction in $\alpha$-helices. J. Mol. Biol. 267:184–197.

[S18] Harper, E. T., and G. D. Rose, 1993. Helix Stop Signals in Proteins and Peptides: The Capping Box. Biochemistry 32:7605–7609.

[S19] Petukhov, M., N. Yumoto, S. Murase, R. Onmura, and S. Yoshikawa, 1996. Factors That Affect the Stabilization of $\alpha$-helices in Short Peptides by a Capping Box. Biochemistry 35:387–397.

[S20] Muñoz, V., F. J. Blanco, and L. Serrano, 1995. The hydrophobic-staple motif and a role for loop-residues in $\alpha$-helix stability and protein folding. Nat. Struct. Mol. Biol. 2:380–385.

[S21] Muñoz, V., and L. Serrano, 1995. Analysis of $i, i + 5$ and $i, i + 8$ Hydrophobic Interactions in a Helical Model Peptide Bearing the Hydrophobic Staple Motif. Biochemistry 34:15301–15306.

[S22] Schellman, C., and R. Jaenicke, 1980. The $\alpha_L$ conformation at the ends of helices., Amsterdam: Elsevier, 53.

[S23] Aurora, R., R. Srinivasan, and G. D. Rose, 1994. Rules for $\alpha$-helix Termination by Glycine. Science 264:1126–1130.

[S24] Viguera, A. R., and L. Serrano, 1995. Experimental Analysis of the Schellman Motif. J. Mol. Biol. 251:150–160.

[S25] Prieto, J., and L. Serrano, 1997. C-capping and Helix Stability: The Pro C-capping Motif. J. Mol. Biol. 274:276–288.

[S26] Lacroix, E., A. R. Viguera, and L. Serrano, 1998. Elucidating the Folding Problem of $\alpha$-helices: Local Motifs, Long-range Electrostatics, Ionic-strength Dependence and Prediction of NMR Parameters. J. Mol. Biol. 284:173–191.
